# Coupling Stickland fermentation with reverse β-oxidation represents a new mode of medium-chain carboxylate production

**DOI:** 10.64898/2026.09.09.750506

**Authors:** Blake G Lindner, Dinesh Kumar Nallasamy, Liam Roberts, Ian M Gois, Christopher E Lawson

**Affiliations:** Department of Chemical Engineering and Applied Chemistry, University of Toronto, Toronto, Ontario M5S 3E5, Canada

## Abstract

Biotechnologies for producing medium-chain carboxylic acids (MCCAs) by chain elongating bacteria (CEB) promise to shift the oleochemical industry away from existing land use practices, vulnerable supply chains, and significant environmental costs. Realizing this potential requires a broader understanding of CEB physiology, which is currently based on lactate and ethanol utilization, despite these bacteria thriving in protein-rich digestive tracts and organic waste bioreactors. Through comparative genomic analysis of known MCCA producers and physiological examination of the recently isolated species *Peptonella octanoica* as a model system, we show that amino acid utilization is a broadly distributed yet largely overlooked trait of many CEB. Among several amino acid-fermenting lineages, a conserved metabolic architecture couples Stickland fermentation with reverse β-oxidation (RβO), enabling MCCA production when acetate or electron-accepting amino acids are limiting. We show how this coupling manages thermodynamic constraints on lactate and amino acid oxidation by driving longer-chain product formation as hydrogen partial pressure increases. We further reveal that amino acid-fermenting CEB are abundant not only in gut microbiomes but also engineered bioreactors, where their novel physiology provides a mechanistic basis for high titer *n*-octanoate production from proteinaceous organic wastes. Beyond biomanufacturing, these findings broaden our understanding of CEB across anaerobic microbiomes in which amino acid fermentation and chain elongation are intersecting ecological strategies.

## Main

Identifying the drivers of metabolite formation among prokaryotes is a central goal of microbial physiology, particularly among anaerobes, which face unique thermodynamic constraints^1^. Anaerobes drive critical processes in human and animal health, food systems, and wide-ranging biotechnologies^2–4^, making mechanistic understanding essential for both fundamental and applied pursuits. Yet genomic and open-culture based discoveries are rapidly outpacing rates of cultivation and characterization leaving the phenotypic potential of most anaerobes unobserved^5–7^. Compounding this issue, fermentative traits are seldom phylogenetically conserved, meaning closely related organisms may diverge in terms of metabolic end products in ways that resist prediction from taxonomy or sequence alone^8,9^.

Chain elongating bacteria (CEB) provide an unusually tractable model for probing the determinants of product diversity among fermentative anaerobes. CEB are a phylogenetically diverse guild united by a core reductive pathway with a key characteristic: variance in terminal carboxylate length which ranges from *n*-butanoate (C_4_) to *n*-octanoate (C_8_). To date, the vast majority of identified CEB produce only short-chain carboxylic acids (SCCAs; C_2_-C_5_) and organisms known to produce medium-chain carboxylic acids (MCCAs; C_6_-C_12_) represent a small and poorly understood minority^10,11^. Yet MCCAs are high-value platform chemicals currently derived from ecologically harmful agricultural practices, for which waste-to-chemical biomanufacturing represents a sustainable alternative^12,13^.

The metabolic foundations of chain elongation were revealed through study of an *n*-hexanoate producer, *Clostridium kluyveri*, which established reverse β-oxidation (RβO) as the core reductive pathway and ethanol as the canonical electron donor^14–18^. Subsequent reactor studies and isolation of additional CEB have revealed lactate as another major substrate and upheld the central role of RβO^19–21^. Since lactate oxidation to pyruvate is thermodynamically unfavourable under many anaerobic conditions, lactate-utilizing CEB encode a flavin-based electron confurcating lactate dehydrogenase (LDH) that couples ferredoxin and lactate oxidation to maintain feasibility across a broad range of physiological conditions^22–24^. Additionally, CEB typically encode both pyruvate:ferredoxin oxidoreductase (PFOR) and pyruvate:formate lyase (PFL), enabling oxidative or non-oxidative acetyl-CoA production from pyruvate. Thus, for studies attempting to elucidate controllers of product length, prevailing models rely entirely on this genetic arrangement and substrate utilization of ethanol, lactate, or carbohydrates^24–26^.

Two interrelated gaps in this framework limit mechanistic understanding and technological development of MCCA biomanufacturing. First, although some CEB inhabit protein-rich anaerobic environments like gastrointestinal systems, fermented food, and organic wastes, the role of amino acid catabolism in redox balancing, energy conservation, and product selectivity in chain elongation remains unknown^27–29^. Following isolation of the non-saccharolytic lactate-utilizing MCCA producer *Peptonella pyruvativorans*^30^, Wallace and colleagues predicted that bacteria deploying the metabolic strategies of *C. kluyveri* “may be more widespread than is generally recognized”^31^. This prediction has been substantiated by the culture record in number but not yet in scientific understanding, in that large numbers of CEB have been isolated, but the vast majority have only been observed producing SCCAs thus hampering efforts to deduce generalizable mechanisms driving MCCA production^9,10,24^. Second, despite the greater industrial value of *n*-octanoate, only three species (*Pseudoramibacter alactolyticus, Megasphaera hexanoica,* and *Megasphaera elsdenii*) have been documented as substantial *n*-octanoate producers prior to our work^24,32^. This scarcity contrasts with productivities of *n*-octanoate and other MCCAs observed in natural microbiomes and engineered open cultures, implying that relevant *n*-octanoate producing CEB remain uncharacterized and the physiologies driving longer product formation incompletely known^33–36^.

To address these gaps, we isolated a collection of chain elongating strains from a continuous open-culture bioreactor fed protein-rich food waste exhibiting robust *n*-octanoate productivity. Screening this collection led to the identification of a novel MCCA-producing CEB capable of producing high titers of *n*-octanoate when fed amino acids or lactate, which we designated *Peptonella octanoica*. Physiological characterization revealed this chain elongator balances redox through both Stickland fermentation and RβO. The novel determinants of its metabolic selectivity towards MCCAs are linked with two departures from canonical lactate-based chain elongation: the absence of PFL and the expression of a non-electron-confurcating LDH. Together, these features impose a thermodynamic constraint on lactate utilization to which *P. octanoica* has two general solutions: 1) when available in sufficient quantities, reduction of an appropriate amino acid, or 2) diversion of greater flux through high cycle number of RβO. This metabolic architecture enables *P. octanoica* to produce high titers of MCCAs (particularly *n*-octanoate) across various media and in batch fermentation of waste feedstocks, demonstrating its biotechnological relevance. The widespread distribution of *Peptonella* species in gastrointestinal environments, combined with the previously unrecognized enrichment of *P. octanoica* in numerous engineered systems, positions this genus as a tractable model for investigating the intersection of amino acid fermentation and chain elongation across both host-associated and biotechnological contexts.

## Results

### The metabolic rules governing MCCA production among Bacillota are incomplete

All known MCCA producers lie within the phylum Bacillota, making it the necessary starting point for deducing generalizable rules about substrate utilization and chain length among CEB. To ground the genetic and phenotypic landscape for our study of terminal product length among organisms within this phylum, we constructed a phylogenetic tree and decorated it with known instances of monocarboxylate production (Figure 1A). To ensure a good amount of daylight reached most branches and attempt a fair display of recognised trait information with minimal taxonomic bias^37^, we selected a single representative genome from each genus within the monophyletic portion of Bacillota (per GTDB r232) containing at least one reference genome accessioned in NCBI’s RefSeq (n=1,107 representative genomes). This allowed us to annotate leaves as either C_4_-C_5_ producers (n=192 leaves; based on a recent multi-domain analysis^8^; Table S1) or known MCCA producers (n=19 leaves; Tables S2-S3).

**Figure 1.**
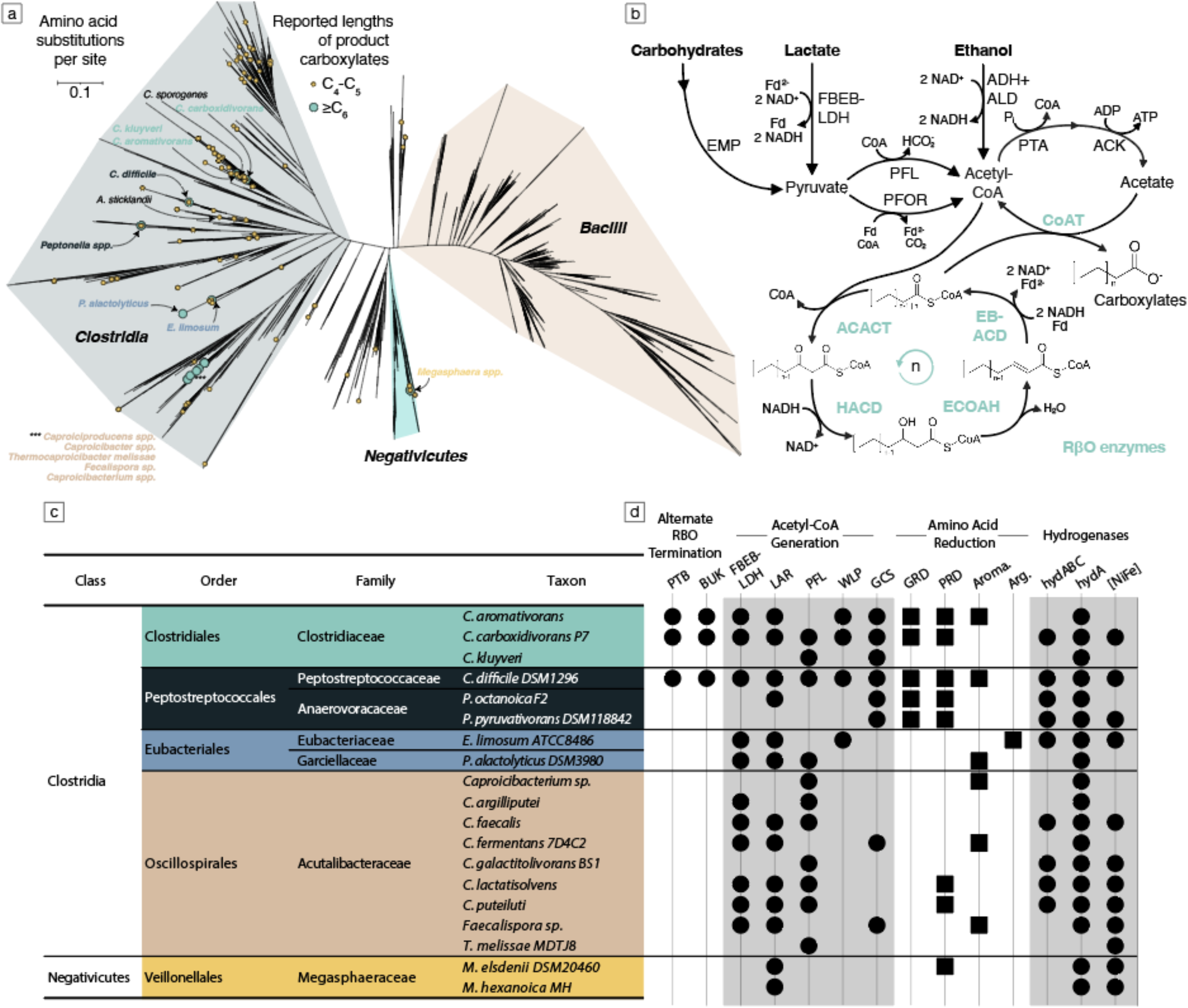
A summary of the state of knowledge regarding C_4_-C_8_ monocarboxylate production among the Bacillota. **Panel A:** Unrooted maximum likelihood tree of concatenated core genes for the phylum drawn from RefSeq entries accessioned in GTDB release 232 and subsampled to genus-level representatives. Shaded regions approximate the boundaries of key taxonomic classes. Coloured circles decorate leaves within which one or more isolates have been experimentally confirmed to produce monocarboxylates of length C_4_ – C_5_ and/or ≥C_6_. Specific clades discussed throughout this work have been labelled for convenience. **Panel B:** The prevailing model of chain elongation with canonical electron donors. **Panel C:** Taxonomic designation of MCCA-producing CEB. **Panel D:** Presence or absence summary of select pathways or genes across CEB reference genomes. Genetic loci are sorted into loose categories (upper labels), and shapes indicate the presence of a particular gene or pathway for the corresponding “Taxon” listed in the panel. Squares are used to emphasize reductive Stickland pathways and circles elsewhere. Horizontal lines extending from Panel C demarcate taxonomic orders. Note that annotation of FBEB-LDH relies only on two electron-transferring flavoproteins (ETFs) and an FAD-dependent LDH homologous to those of *Acetobacterium woodii* without respect to their colocalization. Termination via PTB and BUK is referred to as an alternate mechanism compared to CoAT. Figure abbreviations include Embden-Meyerhof-Parnas pathway (EMP), flavin-based electron bifurcating/confurcating lactate dehydrogenase (FBEB-LDH), pyruvate formate lyase (PFL), pyruvate:ferredoxin oxidoreductase (PFOR), alcohol dehydrogenase (ADH), acetaldehyde dehydrogenase (ALD), phosphate acetyltransferase (PTA), acetate kinase (ACK), coenzyme-A transferase (CoAT), acetyl-CoA C-acetyltransferase (ACACT), 3-hydroxyacyl-CoA dehydrogenase (HACD), enoyl-CoA hydratase (ECOAH), electron bifurcating acyl-CoA dehydrogenase (EB-ACD), adenosine diphosphate (ADP), adenosine triphosphate (ATP), nicotinamide adenine dinucleotide (NADH/NAD^+^), coenzyme-A (CoA), ferredoxin (Fd/Fd^2-^), phosphate butyryltransferase (PTB), butyrate kinase (BUK), nickel-dependent lactate racemase (LAR), Wood-Ljungdahl pathway (WLP), glycine cleavage system (GCS), glycine reductase (GRD), proline reductase (PRD), aromatic amino acid reduction pathway (Aroma.), arginine reduction pathway (Arg.), iron-only flavin-based electron bifurcating/confurcating hydrogenase (hydABC), iron-only ferredoxin-dependent hydrogenase (hydA), and nickel-iron containing hydrogenase ([NiFe]).

We found that, apart from *Bacilli*, nearly every major radiation contains one or more genera observed to produce C_4_ or C_5_ among their major fermentation products. Leaves representing these organisms greatly outnumbered their MCCA-producing counterparts, yet both phenotypes ranged across a similar breadth of the phylogeny. Except for the family *Acutalibacteraceae* (*Caproiciproducens*, *Caproicibacter*, etc.), MCCA production as currently observed is not strongly linked to any taxonomic ranks higher than genus. Though all known MCCA producers rely on RβO for production of *n*-hexanoate or greater, not all C_4_-C_5_ producers must rely on pathways initiated by thiolase-mediated acyl-CoA condensation. Such thiolase-independent production routes are possible via the fermentation of non-canonical electron donors (e.g., leucine, lysine, glutarate, succinate) and reduction of resulting intermediates (e.g., hydroxy acids, enoates) through mechanisms analogous to the latter steps of RβO^38,39^, but which naturally preclude chain elongation.

Although MCCA producers necessarily rely on RβO for production of medium-chain products, we hypothesized that variations in RβO’s flanking catabolic pathways or differences in the usage of electron donors, such as amino acids, could reveal metabolic determinants of terminal product length which would be obscured by a canonical model built around so few reference organisms (Figure 1B). We pursued this question by further interrogating the isolation record via investigation of how well the genetic potential of known MCCA producers aligns with the canonical model of chain elongation. Using reference genomes collected during our tree building efforts (Figure 1C), we began by annotating genomes against multiple major metabolic databases in search of key genes or pathways which would contribute to or detract from conformity with chain elongation canon (Figure 1D).

Our analysis highlighted several discrepancies between chain elongation as a model and the genotypes of organisms confirmed to produce MCCAs. We found that genes predicted to encode flavin-based electron bifurcating/confurcating (FBEB-) LDH and PFL were missing from 8 of the 19 reference genomes. While the former might be conveniently attributed to a lack of lactate utilization (which is well known for some organisms e.g., *C. kluyveri*), the absence of genes encoding PFL from over 40% of MCCA-producing reference genomes cannot be explained by substrate specificity alone. The absence of PFL was accompanied by the presence of some other obvious reductive pathway (i.e., beyond RβO) in every case except *M. hexanoica*. Although some of the observed compensatory reductive pathways were expected – like the Wood-Ljungdahl pathway of known acetogenic CEB – the prevalence of amino acid reduction pathways, specifically genes encoding the selenoprotein reductases of glycine and proline, was unexpected. Together, these genomic observations indicate that our study of the metabolic rules governing MCCA production must extend to include amino acid fermenters in addition to organisms living strictly ethanol-, carbohydrate-, and/or lactate-based lifestyles.

Stickland fermentation has been most frequently described as the pairing of amino acids acting as electron donor and acceptor, with such pairs often yielding acetate, SCCAs, or non-proteinogenic amino acids as products^40–42^. If certain CEB encode amino acid reduction pathways in addition to canonical electron sinks, we hypothesized some lineages may couple oxidative amino acid fermentation pathways directly to MCCA production. To investigate this, we screened for MCCA-producing isolates from a continuously operated open-culture bioreactor fed protein-rich waste, which maintained robust MCCA productivity (0.356 ± 0.048 g L^-1^ day^-1^; 0.121 g MCCA g^-1^ volatile solids yield)^36^. These efforts yielded a novel CEB isolated on lactate-supplemented DSM 104, which we designated *Peptonella octanoica*^43^ after observing large amounts of ammonium and *n*-octanoate in its spent media. The observation of these metabolites seemed to implicate joint operation of RβO and amino acid catabolism, which led us to select *P. octanoica* as a model to investigate how amino acid fermentation and chain elongation cooperate or conflict in the context of MCCA production.

### *Peptonella octanoica* produces high titers of *n*-octanoate both from lactate and amino acids

An early assumption was that *P. octanoica* required canonical electron donors like lactate or ethanol to produce *n*-octanoate. We quickly found this to be incorrect when we observed that *P. octanoica* grown on a modification of its isolation medium (modified DSM 104, hereafter m104) lacking glucose, lactate, or ethanol produced *n*-octanoate (2.7 ± 0.1 mM). This prompted experiments using various m104 formulations informed by high-throughput substrate utilization screening (Figure S1), revealing that *L*-proline and *DL*-lactate drive disparate responses (Figure 2). Growth on m104 with ∼100 mM lactate yielded *n*-octanoate (18.8 ± 1.2 mM) at titers approximately 3– and 10-fold greater than *n*-hexanoate (5.9 ± 0.37 mM) and *n*-butanoate (1.8 ± 0.10 mM), respectively (Figure 2E).

**Figure 2.**
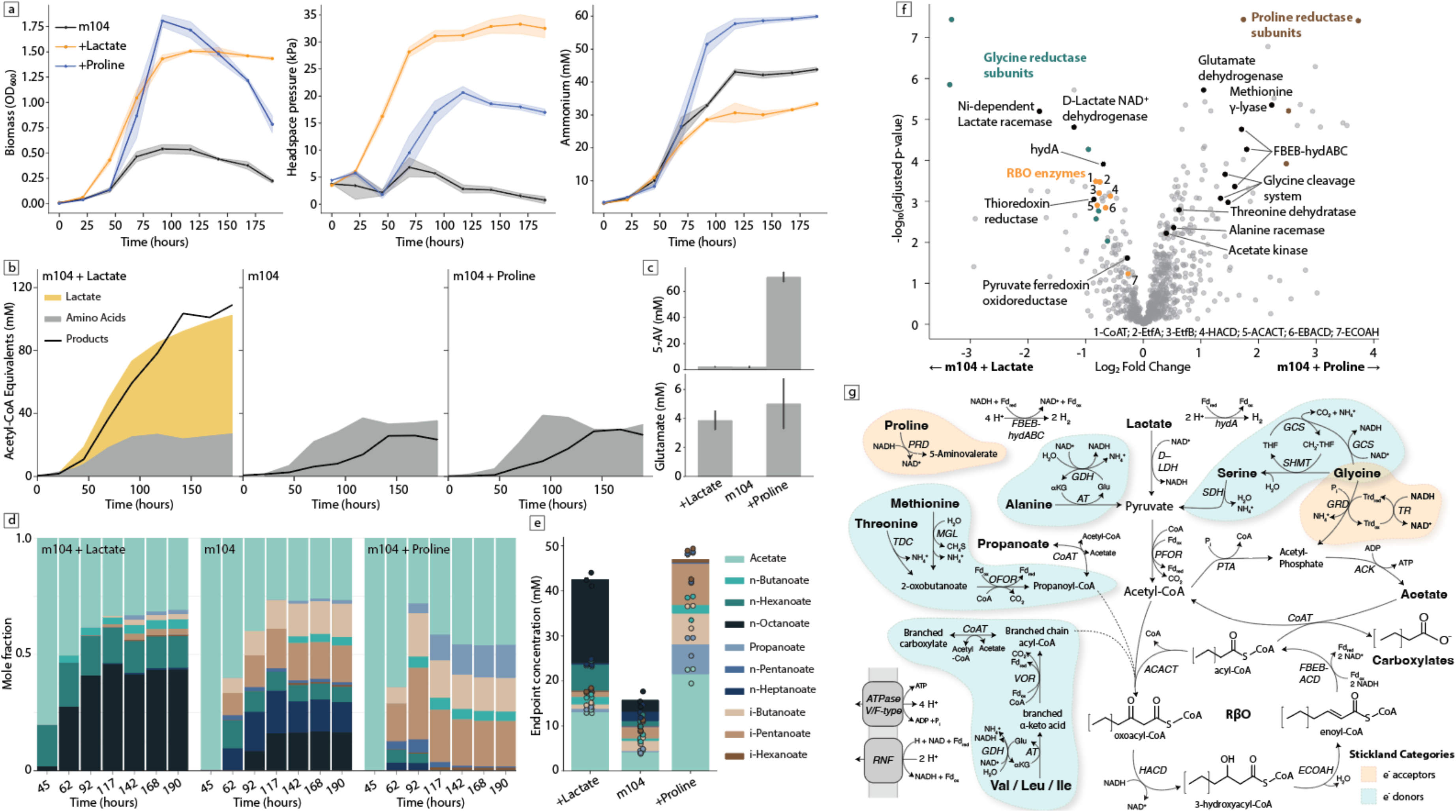
A new model of microbial chain elongation revealed through the metabolism of *P. octanoica*. **Panels A**: Growth, headspace pressure, and ammonium curves across cultivation in different formulations of m104. Solid lines indicate average of biological replicates (n=3) and shaded bounding areas indicate one standard deviation from this mean. **Panels B**: Mass balance in terms of acetyl-CoA equivalents produced in carboxylate products or consumed from lactate or amino acids. Displayed data represents the average of biological replicates. **Panels C**: Fermentation endpoint concentrations of extracellular 5-aminovalerate (top) and glutamate (bottom) across different m104 formulations. **Panels D**: Mean carboxylate proportions (as mole fractions) across fermentation time. Metabolites are sorted from top to bottom with increasing carbon chain length. **Panel E**: Fermentation endpoint carboxylate concentrations. Metabolites are sorted from top to bottom with decreasing carbon chain length. **Panel F**: Differential proteomics between *P. octanoica* cultures grown to mid-exponential phase in m104 amended with lactate or proline. Select proteins are coloured and labeled on the volcano plot with RβO enzymes numbered. **Panel G**: A model of *P. octanoica*’s amino acid and lactate catabolism. Amino acid substrates are drawn within shaded regions indicating their potential for oxidation or reduction. Bolded text indicates substrates and products. Italicized abbreviations indicate enzymes. Abbreviations include proline reductase (PRD), glutamate dehydrogenase (GDH), flavin based electron bifurcating multimeric iron-only hydrogenase (FBEB-hydABC), ferredoxin-dependent monomeric iron-only hydrogenase (hydA), D-lactate dehydrogenase (D-LDH), serine hydroxymethyltransferase (SHMT), serine dehydratase (SDH), glycine cleavage system (GCS), glycine/betaine reductase (GRD), thioredoxin reductase (TR), pyruvate:ferredoxin oxidoreductase (PFOR), coenzyme-A transferase (CoAT), catabolic threonine dehydratase (TDC), methionine gamma-lyase (MGL), 2-oxobutanoate:ferredoxin oxidoreductase (OFOR), 2-ketoisovalerate:ferredoxin oxidoreductase (VOR), aminotransferase (AT), ion-translocating ferredoxin:NAD^+^ oxidoreductase (RNF), thiolase/acetyl-CoA C-acetyltransferase (ACACT), 3-hydroxyacyl-CoA dehydrogenase (HACD), enoyl-CoA hydratase (ECOAH), flavin-based electron bifurcating acyl-CoA dehydrogenase (FBEB-ACD), phosphate acetyltransferase (PTA), acetate kinase (ACK), tetrahydrofolate (THF), and α-ketoglutarate (αKG).

In contrast, growth on ∼90 mM proline generated an approximately equimolar concentration of 5-aminovalerate, increased growth rate, and replaced MCCA production with a diverse set of SCCAs including branched (*i*-butanoate, *i*-pentanoate) and odd-chain (propanoate, *n*-pentanoate) carboxylates (Figure 2A,C,D). Substantial deamination occurred under both conditions, with average exponential phase rates of approximately 1.1 and 0.54 mmol L^-1^ hour^-1^ for proline and lactate conditions, respectively. This deamination was the result of extensive amino acid utilization, including primarily alanine, leucine, and valine (between 70-100%; Table S5). Consequently, when expressed as acetyl-CoA equivalents, product formation could not be accounted for by lactate utilization alone. Inference of amino acid catabolism’s contribution to acetyl-CoA units based on ammonium accumulation was required to balance products and reactants (Figure 2B). Additional m104 experiments confirmed that *P. octanoica* also utilizes SCCAs as co-substrates for RβO (Figure S2).

Our inability to balance carboxylate production solely with lactate utilization prompted further genomic and proteomic investigation. These efforts highlighted connections between three major metabolic systems within *P. octanoica*: RβO, amino acid oxidation pathways, and Stickland reductases for proline and glycine (Figure 2F). Under proline conditions, the glycine cleavage system and proline reductase were differentially expressed. Under lactate conditions, RβO enzymes and a glycine reductase were differentially expressed. Across both conditions, amino acid oxidation flexibly paired with either RβO or Stickland reduction via constitutively expressed oxidoreductases paralogous to PFOR (i.e., OFOR, VOR; Figure 2G; Tables S6-S7). These enzymes generate acyl-CoA intermediates from ketoacids produced by amino acid trans/deamination, enabling entry into RβO or release to drive substrate-level phosphorylation. Under proline-fed conditions, high electron acceptor availability enabled redox balance with minimal RβO flux, resulting in higher growth rates at the expense of MCCA production (Figure 2A,D).

As described above, we observed a lack of genes encoding the multimeric electron-confurcating LDH and PFL associated with model chain elongators and other non-respiring lactate-oxidizing anaerobes^13,22,44^ (Figure 1B,D). In these experiments, consistent with the lack of PFL, extracellular formate was not detected under any condition described here or below. We observed statistically significant upregulation of an electron confurcating iron-only hydrogenase (hydABC) and a monomeric iron-only hydrogenase (hydA) in the presence of proline and lactate, respectively. Additionally, a nickel-dependent lactate racemase and a putative *D*-lactate dehydrogenase were differentially expressed on lactate (Table S8; Figure S3). Expression data, metabolic reconstruction, and annotation information are further detailed in Supporting Information (Tables S6-S7).

To resolve how *P. octanoica*’s dual amino acid-fermenting and chain-elongating lifestyle compares to a closely related CEB and a model amino acid fermenter, we cultivated *P. octanoica*, *P. pyruvativorans*, and *Clostridium sporogenes* in 10% m104 supplemented with casamino acids (20 g L^-1^) and lactate or ethanol provided as additional electron donors (Figure 3A). All strains depleted proline and glycine completely and produced comparable quantities of 5-aminovalerate regardless of electron donors (Figure 3B.D). *Peptonella* species produced MCCAs across all media compositions, while *C. sporogenes* yielded no carboxylates longer than C_4_ under any conditions tested (Figure 3C). *C. sporogenes* drew on a broader range of amino acids than either *Peptonella* species, including much greater consumption of aromatic amino acids (Figure 3D). Although *C. sporogenes* encodes the enzymes for RβO^45^, we only observed its operation in the presence of lactate for *n*-butanoate production which contrasts with how RβO appears constitutively integrated with amino acid catabolism among *P. octanoica* and *P. pyruvativorans*.

**Figure 3.**
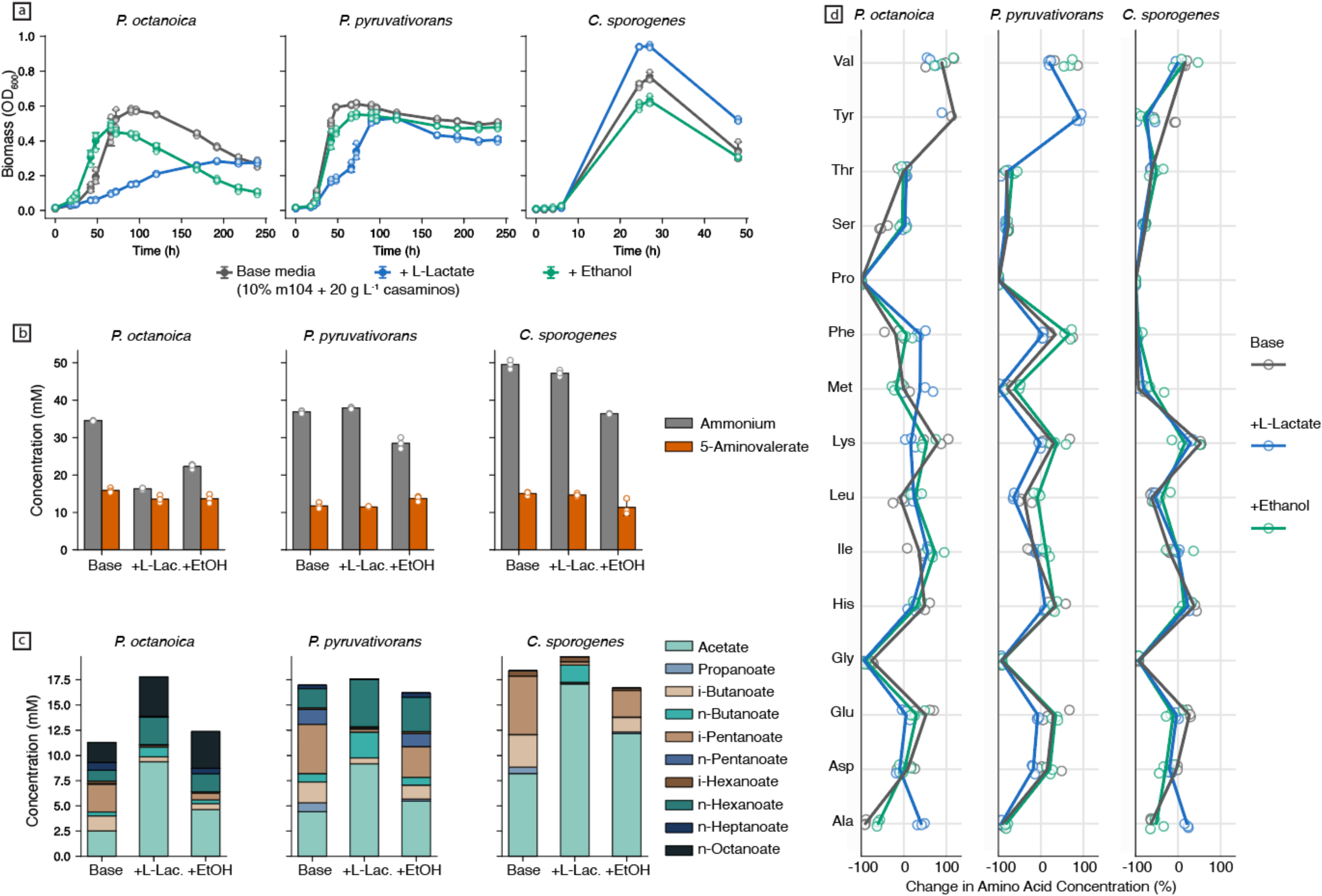
Species of *Peptonella* vary in amino acid utilization strategies compared to *C. sporogenes*, an SCCA-producing Stickland fermenter. **Panels A**: Growth curves across three formulations of 10% m104 with 20 g L^-1^ casamino acids with either added lactate or ethanol. Solid lines represent the average of biological replicates (n=3) and error bars correspond to one standard deviation with individual replicates overlaid. **Panels B**: Deamination and 5-aminovalerate formation at experimental endpoints. Bar heights represent averages of biological replicates (n=3). **Panels C**: Carboxylate formation at experimental endpoints. Stacked bar heights are averages of biological replicates (n=3). **Panels D**: Change in extracellular amino acid concentrations between experimental endpoint and initial conditions. Biological replicates (n=3) are plotted as individual points and solid vertical lines connect the average of these replicates. Solid lines tracing mean percent change are not shown for species and amino acids combinations with observed changes exceeding 150%. Colours correspond to media formulations. Abbreviations include L-lactate (L-Lac.) and ethanol (EtOH).

To understand this divergence in RβO output between phylogenetically and/or functionally similar organisms, we turned to recent work proposing mechanisms that control product formation differences within the pathway. Comparative analysis of *M. hexanoica* and an *n*-butanoate-producer has shown that the thiolase (ACACT) enzyme of RβO is a key molecular determinant of product length via a mechanism involving steric selectivity at the acyl-CoA binding pocket^25^. Similar efforts comparing the activity of the CoA transferase (CoAT) of *Pseudoramibacter alactolyticus* to that of an SCCA-producer revealed an analogous molecular mechanism for product termination^24^. Taken together, these results reveal how both enzymes act as joint coordinators for the entrance and exit of acyl-CoA molecules from RβO. Consequently, we next compared the genes encoding the RβO pathway of *P. octanoica* (as identified in Figure 2F) to *P. pyruvativorans*, other relatives, and MCCA producers.

### Coding differences in RβO enzymes sustain variations in substrate utilization and product formation among amino acid-fermenting CEB

We found that each RβO enzyme differentially expressed in the presence of lactate is co-localized in an operon-like arrangement on the genome of *P. octanoica* (Figure 4A). Consistent with its observed constitutive expression, ECOAH is encoded adjacent to and on the opposite strand from this gene cluster. This arrangement, particularly the insertion of a CoAT-encoding gene into the RβO cluster, is uncommon^46^. In the intergenic region between ECOAH and HACD, we observed two operator sites for redox-sensitive repressor Rex. Based on these observations, we investigated genomic loci among three groups: other MCCA producers, the family *Anaerovoracaceae*, and the genus *Peptonella*. When examining the broader scope of all MCCA producers, the adjacency of RβO enzymes seems to roughly obey taxonomic boundaries i.e., gene adjacency was most similar among MCCA producers of the same order (Tables S9-S10). Amino acid local alignment identity was consistently above 50% for ECOAH, HACD, and ACACT and 60% for EB-ACD, except for the EB-ACD sequences of *Megasphaera,* which appeared more divergent (Figure S4).

**Figure 4.**
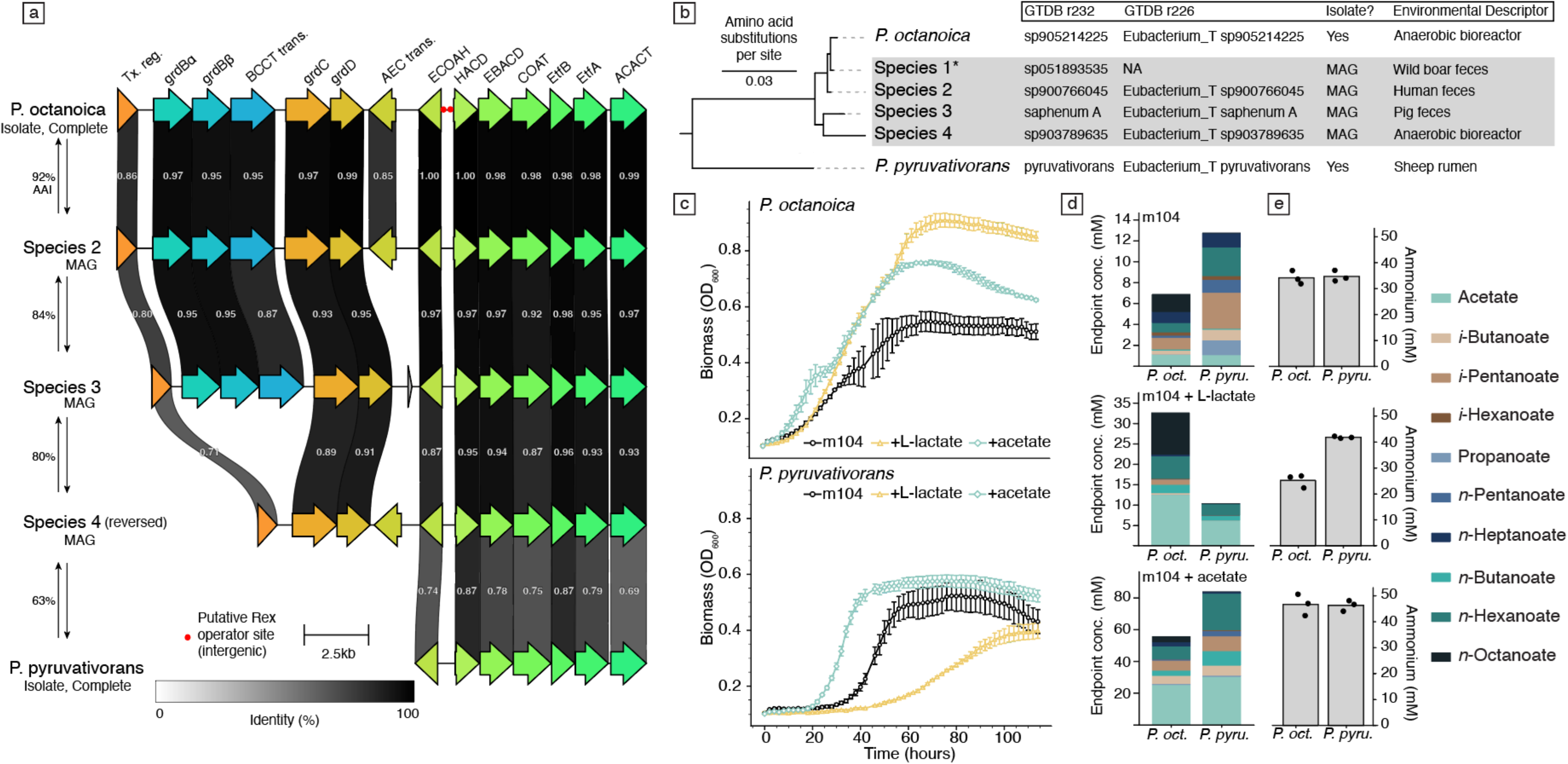
Distribution and similarity of the genetic loci enabling RβO across the genus *Peptonella*. **Panel A**: A conserved operon and its context across *Peptonella*. Coding sequences are indicated with coloured arrows. Ribbons connecting coding sequences are labelled with pairwise amino acid identities. Darker shades illustrate higher identities. Paired arrows on the left side are labelled with average amino acid identity (AAI) between genome pairs. Row order is arranged to reflect the phylogenetic relationship displayed in the next panel. **Panel B**: The phylogenetic relationship between known species-level clades of *Peptonella* and their origins as isolates or metagenome-assembled genomes (MAGs). Phylogenetic distance corresponds to a concatenated core gene tree. Leaf labels from recent releases of GTDB (232 and 226) are also indicated. Species 1-4 reflect species-level clades which have yet to be isolated or named but have been observed in metagenomic datasets. Environmental descriptors correspond to the source of isolation (for isolates) or the origin of the metagenomic dataset producing the MAG selected as species representative in GTDB. **Panels C**: Averages of growth curves from *Peptonella* isolates grown in m104. Error bars indicate standard deviations of biological replicates (n=3). **Panels D**: Average metabolite concentration at fermentation endpoints of *Peptonella* isolates across different m104 compositions. Initial concentrations for added substrates were approximately 50 mM and 100 mM for m104 with acetate and m104 with L-lactate, respectively. **Panels E**: Average endpoint ammonium concentrations with plotted biological replicates corresponding the media compositions in the labels of Panel D. *Species 1 is not included in illustration of the RβO gene cluster shown in Panel A because its gene cluster exists on a short contig with edges adjacent to ECOAH and ACACT. Abbreviations include transcriptional regulatory (Tx. reg.), glycine/betaine reductase (grd), betaine transporter (BCCT trans.), and auxin efflux channel-like transporter (AEC trans.).

At the family level, homologs to *P. octanoica*’s RβO genes were observed in most *Anaerovoracaceae* reference genomes. Despite the closer phylogenetic relationship, we found amino acid identities at the family level were essentially comparable to those between *P. octanoica* and other MCCA producers reported above, except for modestly elevated ECOAH identities (Figure S5). In contrast, alignment between *P. octanoica* and other members of its genus revealed that a syntenically intact RβO gene cluster is a universally shared feature (Figure 4A,B). Further, most of *Peptonella*’s species-level clades encode RβO operons which are flanked by loci dedicated to glycine reduction with *P. pyruvativorans* as a notable exception – though the subunits of this reductase are encoded elsewhere on its genome. When comparing the similarity of RβO genes across this syntenic region, we observed amino acid identities for all RβO genes that exceeded their pairwise genome-wide average amino acid identities (AAI). Within this conserved RβO gene cluster, we found that CoAT was consistently the most divergent gene across the clade, followed by ACACT. This observation is consistent with the emerging notion in the literature that both enzymes control RβO product length.

We hypothesized that divergence between initiating and terminating RβO enzymes, combined with an organism’s broader physiology, drives meaningful differences in the metabolite profiles of *Peptonella* species. To probe this theory, we compared growth and extracellular metabolites of *P. octanoica* and *P. pyruvativorans* in m104 with added L-lactate or acetate (Figure 4C). In basal m104, both organisms produced similar quantities of ammonium, but their carboxylate product profiles differed (Figure 4D,E). *P. pyruvativorans* produced substantially greater amounts of branched– and odd-chain carboxylates, while *P. octanoica* yielded a greater proportion of even-chain products, inclusive of *n*-octanoate. Deamination quantities across basal and L-lactate containing m104 were consistent for *P. pyruvativorans,* whereas the presence of lactate decreased *P. octanoica*’s amino acid utilization. In contrast, both species exhibited increased amino acid utilization in the presence of acetate – a known response among Stickland fermenters^40^. Despite *P. pyruvativorans* encoding a high-identity homolog of *P. octanoica*’s D-LDH, we observed substantial growth repression and depression of metabolite formation in the presence of lactate, which we found was due to the absence of lactate racemase and not an inability to oxidize lactate (Figure 1D).

Next, we investigated the growth and metabolite formation of other MCCA producers in the presence of amino acids to determine the generalizability of pairing amino acid catabolism and RβO typified by *Peptonella*. Based on the genomic capacity for amino acid catabolism observed in *Megasphaera* (Figure 2), we examined growth of *M. hexanoica* across m104 formulations. In basal media, *M. hexanoica* supported minimal growth (0.19 ± 0.01 max. OD_600_) and yielded significantly less ammonium (9.6 ± 0.77 mM NH_4_^+^) than *Peptonella* grown under identical conditions (Figure S6). Supplementation with L-lactate or acetate stimulated growth and deamination, with acetate producing a larger effect (0.72 ± 0.13 max. OD_600_; 34.6 ± 3.28 mM NH_4_^+^). MCCA production was observed across conditions, though *n*-octanoate appeared only when lactate or acetate were supplied; *n*-heptanoate dominated in basal medium while *n*-hexanoate was most abundant in lactate– and acetate-supplemented conditions. Additionally, when cultivating the carbohydrate-fermenting MCCA producer *C. fermentans* in m104 formulations with glucose and L-lactate, we did not observe amino acid utilization. In contrast, product carboxylate profiles for *M. hexanoica* and *P. octanoica* were broadly consistent across these conditions, including in basal media which lacked canonical electron donors (Figure S7). Overall, these results demonstrate that integration of amino acid catabolism and RβO are not exclusive to the MCCA producers of *Peptonella* and that phylogenetically distant *Megasphaera* can achieve a similar metabolic pairing, though with greater reliance on co-substrates such as acetate.

### RβO sustains anaerobic lactate oxidation and amino acid utilization in the absence of proline

To determine how Stickland reduction interacts with RβO, we grew *P. octanoica* on 10% m104 with ∼100 mM DL-lactate and proline across a range of molar ratios (Figure 5A). Growth rates increased directly with higher proline:lactate ratios, and near-complete lactate utilization (99.7 ± 0.03%) was achieved at ∼150 mM proline (3:2; Figure 5B). Rising proline concentrations progressively shifted metabolite production toward SCCAs. At an equimolar ratio of proline and lactate, *n*-octanoate production ceased entirely, with the majority of substrate converted to acetate and nominal titers of *n*-butanoate and *n*-hexanoate (Figure 5B,C). Increased Stickland reduction supported by increasing concentrations of proline enabled faster growth rates driven by less reliance on the enzymatically expensive operation of RβO^13,24^ needed for medium-chain product formation (Figure 5G). Proline was completely consumed in all conditions tested, resulting in equimolar 5-aminovalerate (Figure 5H).

**Figure 5.**
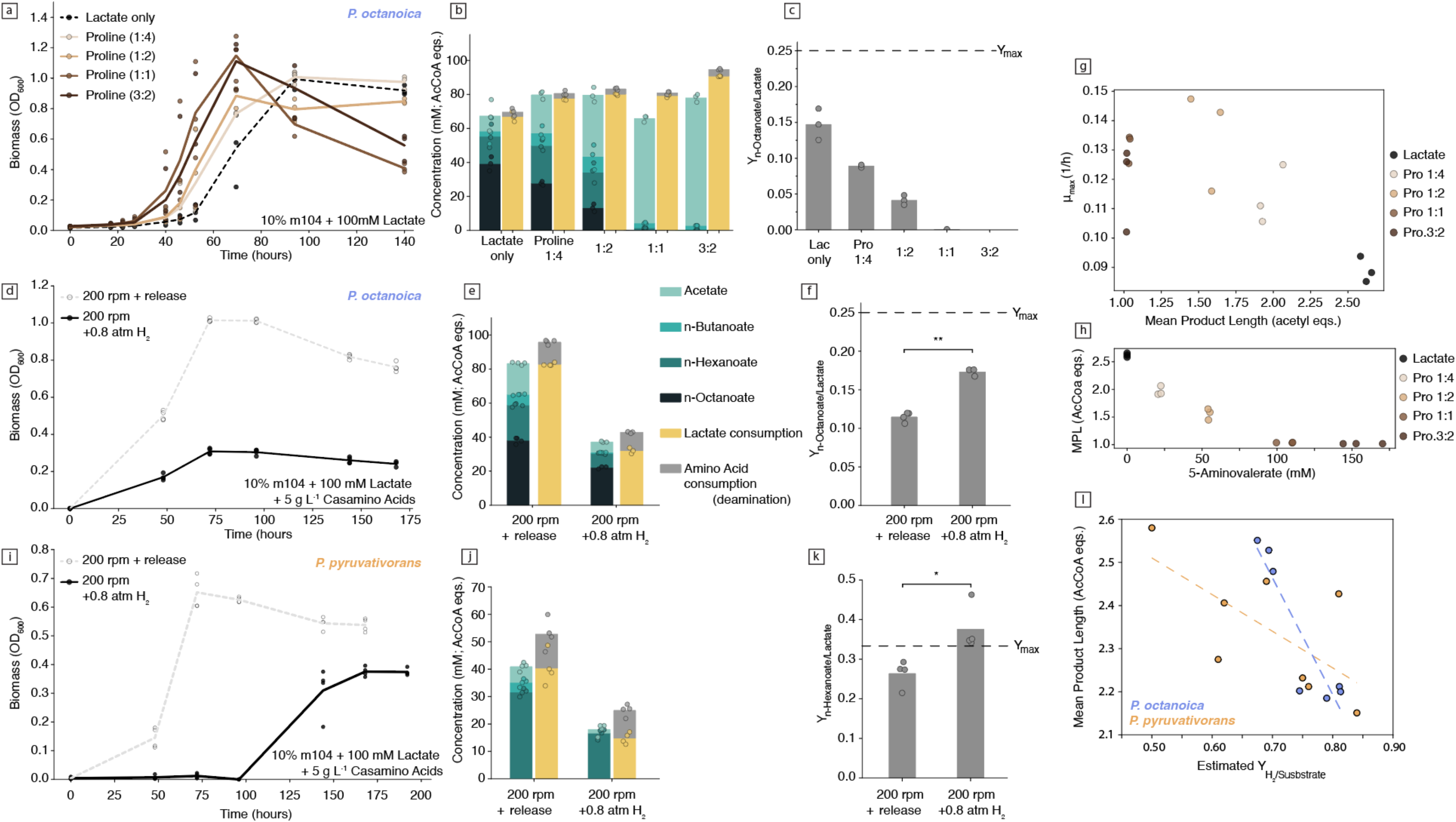
The addition of exogeneous H_2_ constrains lactate utilization which *Peptonella* can partially overcome via production of longer chain products. **Panel A**: Growth curves of *P. octanoica* in 10% m104 with 100 mM DL-lactate and varying amounts of proline. Solid lines represent the mean of the plotted biological replicates (n=3). **Panel B**: Product and substrate bar plots indicating monocarboxylate production (left of pair) and lactate or amino acid utilization (right of pair). Bar heights represent the average of biological replicates. The x-axis is labelled with proline-to-lactate ratios. **Panel C**: Molar yield of *n*-octanoate per mol of consumed lactate for *P. octanoica* grown with varying concentrations of proline and lactate. Bar heights represent the average of biological replicates and maximum theoretical n-octanoate yield per mol of lactate (Y_max_) is labeled with a horizontal dashed line (0.25). **Panel D**: Growth curves of *P. octanoica* in 10% m104 with 100 mM DL-lactate and 5 g L^-1^ casamino acids grown with or without elevated H_2_ partial pressure. Solid lines represent the mean of the plotted biological replicates (n=3 and 4 for pressurized and released conditions, respectively). **Panel E**: Product and substrate bar plots from the conditions for the growth curves displayed in Panel D. Bar heights represent the average of biological replicates. **Panel F**: Molar yield of *n*-octanoate per mol of consumed lactate for *P. octanoica* grown under differing H_2_ partial pressures. Bar heights represent the average of biological replicates and maximum theoretical n-octanoate yield per mol of lactate (Y_max_) is labeled with a horizontal dashed line (0.25). **Panel G**: Maximum observed growth rates (μ_max_) and mean product length (in C_2_ units) for biological replicates of *P. octanoica* grown across various proline-to-lactate ratios. **Panel H**: Mean product length (C_2_ units) plotted against extracellular 5-aminovalerate concentration across proline-to-lactate ratios. **Panel I**: Growth curves of *P. pyruvativorans* in 10% m104 with ~100 mM DL-lactate and 5 g L^-1^ casamino acids grown with or without elevated H_2_ partial pressure. Solid lines represent the mean of the plotted biological replicates (n=4). **Panel J**: Product and substrate bar plots from the conditions for the growth curves displayed in Panel I. Bar heights represent the average of biological replicates. **Panel K**: Molar yield of *n*-hexanoate per mol of consumed lactate for *P. pyruvativorans* grown under differing H_2_ partial pressures. Bar heights represent the average of biological replicates and maximum theoretical *n*-hexanoate yield per mol of lactate (Y_max_) is labeled with a horizontal dashed line (0.33). Due to the lack of racemase activity, amino acid utilization is proportionally larger in the case of *P. pyruvativorans*’ enabling n-hexanoate molar yields greater than possible given the lactate utilization observed. **Panel L**: Mean product length (C_2_ units) plotted against estimated molar yield of H_2_ normalized to combined substrate utilization (lactate and amino acids) from growth experiments in Panels D and I. Hydrogen yield is based on the model introduced in Figure 3, specifically via redox balances solved for flux through a ferredoxin-dependent hydrogenase as the sole unknown. Plotted biological replicates are coloured according to species and fitted with a linear trend as displayed. Individual points in panels B, E, F, J, and K are displayed with minor horizontal jitter to reduce overlap. Statistical significance between conditions in panels F and K was assessed with Welch’s t-test and indicated as *p* < 0.05 (*) and *p* < 0.01 (**\*\***). Figure abbreviations include mean product length (MPL) and acetyl-CoA equivalents (AcCoA eqs.).

Consequently, we theorized that the genus*’* product preference is a response to a fundamental thermodynamic problem: lactate oxidation under anaerobic conditions. The lack of an electron-confurcating LDH means that the organisms’ metabolism must contend with the narrow feasibility of NAD-dependent lactate oxidation. Further exaggerating this issue is the lack of a non-oxidative route for producing acetyl-CoA from pyruvate, which results in invariable production of reduced ferredoxin by PFOR. We hypothesized that *Peptonella*’s bias towards MCCA production is an adaptative response to thermodynamic constraints requiring maintenance of an intracellular state ensuring the feasibility of both lactate oxidation and the forward action of glutamate dehydrogenase (GDH) essential for oxidation of certain amino acids (Figure S8; Table S11). That is, flux through RβO to products of greater length more efficiently disposes of electrons on product carboxylates which drives the re-oxidation of NADH needed to sustain these NAD-dependent reactions without the need for alternative acceptors. This type of adaptation is consistent with how lactate-to-pyruvate ratios have been found indicative of intracellular NADH/NAD^+^ maintenance in other systems^47^ and may result in lower lactate affinities among *Peptonella* when compared to their electron-confurcating lactate-utilizing peers.

The turnover of pyruvate to acetyl-CoA needed to initiate RβO also requires re-oxidation of ferredoxins carrying low potential electrons for which proton translocation via the RNF complex or hydrogen evolution are the two main solutions (Figure 3G). Thus, we tested our hypothesis by inhibiting hydrogenase-mediated ferredoxin oxidation through artificially elevating H_2_ partial pressure (max pH_2_ = 0.8 atm) in *Peptonella* cultures grown on 10% m104 with ∼100mM lactate and minimal casamino acids (5 g L^-1^). To simulate semi-constant pH_2_ we employed vigorous shaking and undertook daily sparging and re-pressurization with fresh gas mix in experimental samples or the release of accumulated gases from controls. We found *Peptonella* growth rates and maximum OD_600_ declined with increasing pH_2_ (Figure 5D,I). Specifically for *P. octanoica*, lactate utilization was inversely correlated with pH_2_ ranging from 80.8 ± 0.7% to 31.3 ± 1.3% (Figure 5E; Figure S9). Average product length increased and *n*-octanoate yield reached 0.173 ± 0.004 mol C_8_ mol^-1^ lactate consumed under high pH_2_ (Figure 5F).

For *P. pyruvativorans*, given its lack of lactate racemase, we observed less utilization overall but similar values when factoring in the availability of D-lactate within the provided racemic mixture. That is, *P. pyruvativorans* consumed 73.2 ± 3.6% of D-lactate compared to only 29.4 ± 2.5% under high pH_2_ conditions. As with *P. octanoica*, the addition of hydrogen shifted product length significantly, and for *P. pyruvativorans* this led to the complete erasure of *n*-butanoate production yet did not stimulate *n*-octanoate production, rather flux through RβO was routed entirely to *n*-hexanoate (Figure 5J). As a result, the observed *n*-hexanoate yield based on lactate consumption rose from 0.264 ± 0.03 to 0.376 ± 0.05 mol C_6_ mol^-1^ lactate, exceeding the maximum yield due to a higher proportion of amino acid utilization (Figure 5K).

For both organisms, the observed increase in MCCA production corresponded with a general decrease in predicted flux through ferredoxin-dependent hydrogenases (Figure 5L). We found these trends corroborated stoichiometries predicted via the model we introduced above for the growth of *Peptonella* on lactate in the absence of acetate. That is, regardless of hydrogenase type, longer chain products drive progressively lower molar hydrogen yields per mole of substrate utilized, thus enabling continued growth under rising pH_2_ (Table S12). Of course, this metabolic flexibility represents a strategy with a limit, as lactate concentrations decrease and/or pH_2_ elevates, lactate utilization necessarily comes to a halt due to thermodynamic infeasibility. Next, we sought to demonstrate how this metabolic architecture performs outside of controlled media compositions to confirm whether *Peptonella* could sustain high titer MCCA production directly from unmodified waste feedstocks.

### Amino acid fermenting CEB are industrially relevant chassis for waste-to-chemical biomanufacturing

To inform efforts valorizing real organic wastes and to map the ecological contexts supporting *Peptonella*, we collected publicly available metagenomic datasets with detectable abundances of isolated genus members and manually collated relevant metadata from associated SRA accessions^48^ (Table S13). We found that *P. octanoica* and *P. pyruvativorans* have substantial differences in habitat preference and range. Both species inhabit the human gut with relatively equal abundances, but *P. pyruvativorans* appears to be higher ranked in most host-associated microbiomes compared to *P. octanoica* (Figure 6A). For metagenomic datasets containing *P. octanoica* signal, we independently mapped reads to assess the level of strain similarity between in situ populations and the reference genome of *P. octanoica* F2 (Figure 6B). Among bioreactor-associated metagenomes, higher strain identities were linked with reactor feed type, and populations in metagenomes from reactors fed dairy residuals exhibited the highest strain-level similarities (Figure 6C).

**Figure 6.**
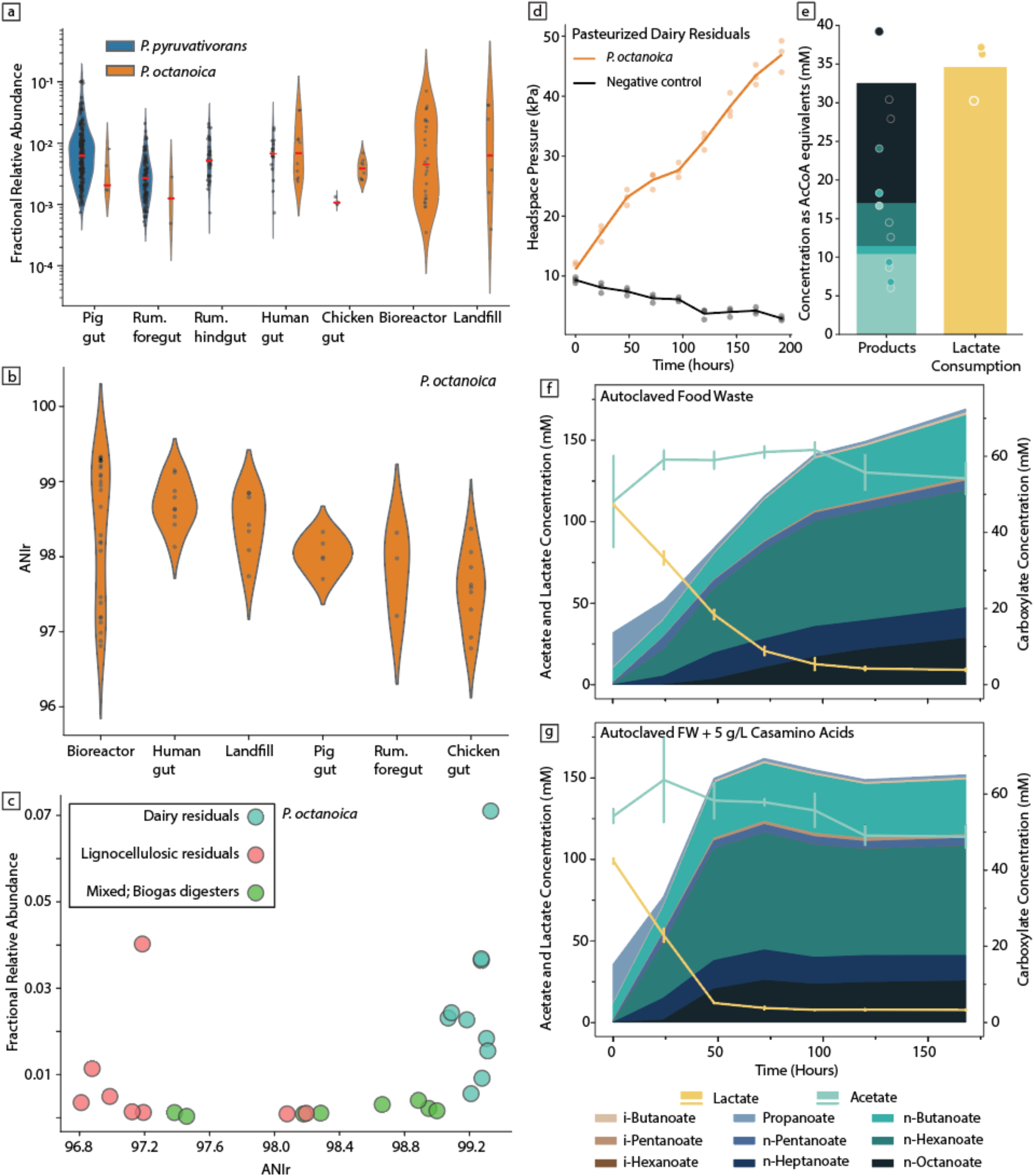
*Peptonella* ecology and applications in waste-to-chemical biotechnology. **Panel A:** Habitat-specific distribution of *P. octanoica* and *P. pyruvativorans* across publicly available metagenomic datasets collected via Sandpiper. The relative abundances displayed on the y-axis have been log_10_ transformed, violin plot areas are normalized individually by habitat, and horizontal red markings indicate medians of relative abundance distributions. **Panel B**: Strain-level identities between metagenomic populations and *P. octanoica* reference genome versus habitat type. Points represent a single metagenomic dataset. **Panel C**: Relative abundance versus strain-level identity within bioreactor metagenomes according to feed type. **Panel D:** Headspace pressure changes over time in pasteurized acid whey inoculated with *P. octanoica* against negative controls. **Panel E**: Endpoint metabolite profiles of *P. octanoica* inoculated acid whey. The metabolite concentrations (y-axis) have been transformed to acetyl-CoA equivalents to better illustrate the balance between substrate and product. **Panel F**: Time series metabolites of *P. octanoica* inoculated into autoclaved food waste (source-separated organics). The primary y-axis indicates acetate and lactate concentrations (lines) and the secondary y-axis corresponds to product monocarboxylate concentrations (areas). Product concentrations remain untransformed (i.e., not as acetyl-CoA equivalents). Error bars indicate one standard deviation among biological replicates (n=3). **Panel G**: *P. octanoica* inoculated autoclaved food waste supplemented with minimal casamino acids (5 g/L). Axis, error bars, and unit conventions follow those of Panel F. Uninoculated negative controls for autoclaved food waste fermentation are in the supporting information (SI 21). Individual points in panels A and E are displayed with minor horizontal jitter to reduce overlap. Figure abbreviations include ruminant (rum.), average nucleotide identity of mapped reads (ANIr), and acetyl-CoA (AcCoA).

Given the high abundance of *P. octanoica* populations inhabiting bioreactors fed dairy and its isolation from source-separated organics (food waste), we investigated its performance as an engine for waste-to-chemical MCCA production (Figure S10). We inoculated pasteurized acid whey and autoclaved food waste in batch fermentation formats and measured the formation of product monocarboxylates relative to uninoculated controls (Figure 6D; Table S15). Both wastes supported robust growth as measured by headspace pressure (Figure 6D) and product formation (Figure 6E,F), including *n*-octanoate. The addition of casamino acids to food waste accelerated lactate utilization without substantial modification of product profiles (Figure 6G).

## Discussion

One third of all global food production is wasted^49,50^, disproportionately contributing to half of all food-related greenhouse gas emissions^51^. Chain elongating biotechnologies could redirect more than a billion tonnes of this organic waste into chemical feedstocks, yet established models have not accounted for amino acids (20-50% of food waste by dry weight^52^) as substrates. Our work fundamentally expands the model of microbial chain elongation in three ways.

First, we demonstrate that amino acids serve as primary substrates from which CEB can form MCCAs. Pyruvate-yielding amino acids such as alanine and serine are the clearest examples, with these and other amino acids oxidized to acyl-CoA intermediates that either drive substrate-level phosphorylation via oxidative Stickland reactions or enter RβO for elongation. The latter is mechanistically divergent from reductive amino acid pathways which circumvent Claisen condensation – such as those used by *C. sporogenes* to reduce leucine, lysine, glutamate, and phenylalanine – instead funneling carbon into longer products; this is likely the route by which *P. octanoica* generates trace *i*-heptanoate in its spent media^43^. Utilizing amino acids in this manner is distinct not only from the view of Stickland fermentation informed by study of model amino acid fermenters^39–41,53–55^, but also from the canonical view of chain elongation. The emergence of MCCA production as an adaptive metabolic response is analogous to the acetogenic flexibility documented in some of those same model amino acid fermenters, namely *A. sticklandii*, *C. difficile,* and *E. limosum*^41,56,57^.

Second, we show how competing reductive pathways shape carboxylate profiles in amino acid-fermenting MCCA producers. Reductive Stickland pathways proved surprisingly prevalent among MCCA producers, with genes encoding these pathways present in more than half of the reference genomes analyzed. In *P. octanoica*, elevated proline concentrations shifted product formation towards SCCAs, approaching nearly complete acetate production at extreme proline:lactate ratios. This product shift enables a change in the organism’s energy conservation strategy, decreasing reliance on IMF generated in tandem with RβO and increasing SLP through acetyl-CoA release to generate greater ATP yield. A further dimension of energy conservation is linked to lower H_2_ partial pressure, which ensures the thermodynamic feasibility of electron-confurcating hydrogenase (HydABC);^58^ consistent with this, HydABC was differentially expressed under proline-fed conditions relative to lactate-fed conditions, where the simpler ferredoxin-dependent hydrogenase (HydA) was expressed instead. Proteomic data confirmed reciprocal regulation of glycine pathways^59^ and proline reductase, consistent with regulatory networks described in *C. difficile*^39,55,60,61^ and highlighting how amino acid availability modulates the physiology of amino acid fermenting CEB.

While some amino acid fermenters rely on syntrophic partnerships with H_2_-consuming populations to continue fermentation when Stickland acceptors are limiting, MCCA production may have arisen as an alternative metabolic strategy – one that is consistent with our observation of MCCA production by amino acid fermenters in a methanogenesis-arrested system. Previous work identified populations of *Anaerovoracaceae* closely related to *Peptonella* as predictors of MCCA production and selectivity in H_2_ co-feeding experiments under methanogen inhibition^62^. The framework described here provides a mechanistic basis for this observation. By directing reducing equivalents to longer and more reduced carboxylates, amino acid-fermenting CEB can minimize hydrogen production, at the cost of growth rate, to sustain their metabolism in the absence of syntrophic partners. More broadly, the responsiveness of amino acid-fermenting MCCA producers to H_2_ partial pressure suggests that hydrogen-consuming community members may shape CEB product profiles in natural and engineered settings with direct implications for the design of chain elongation biotechnologies.

Third, we demonstrate that product chain length is a tunable parameter modulating the feasibility of thermodynamically challenging oxidative reactions. Elevated H_2_ partial pressure inhibited *Peptonella* growth and decreased lactate utilization while simultaneously shifting product profiles towards longer products consistent with higher RβO cycles serving as a flexible redox balancing mechanism when hydrogenase activity is constrained. Supporting this interpretation, our metabolic model for *Peptonella* corroborates that longer products drive decreasing hydrogen yields per mole of substrate across both species. We hypothesize that this mechanism extends to facilitating GDH which could be the source and driver of the characteristic MCCA formation observed during the growth of *Peptonella* on peptone without added lactate or proline. Deviations in lactate oxidation among MCCA-producers is supported by our comparative genomic analysis where we found considerable ambiguity in the annotations of LDH-encoding genes. *Megasphaera* is a key genus with known amino acid utilizing tendencies which appear to lack obvious electron-confurcating LDH and previous work with an enzyme of *M. elsdenii*^63^ suggests that lactate oxidation may directly couple with RβO. Our work emphasizes the need for more comprehensive review of the substrate utilization patterns among MCCA producers to address these knowledge gaps and further inform our understanding of CEB physiology.

Additionally, this work advances our understanding of amino acid fermenters and their adaptation to various niches. Specifically, organic acid stress represents an additional selective pressure favouring MCCA production from lactate-rich environments. *Peptonella* tolerates pH as low as 4.8 facing high acetate and lactate concentrations (50-100 mM) – conditions challenging for many clostridial amino acid fermenters. Conversion of lactate to less acidic longer-chain products offer a physiological advantage when balanced against the increasing antimicrobial activity of longer MCCAs. This balance may be accomplished via niche differentiation within *Peptonella* itself. For example, *P. octanoica* is a DL-lactate utilizing *n*-octanoate producer while *P. pyruvativorans* utilizes a broader range of amino acids but is restricted to stereoselective lactate utilization and the production of *n*-hexanoate. Among these two species, *P. octanoica* was observed with high prevalence in engineered lactate-rich settings whereas *P. pyruvativorans* was predominantly found in host-associated environments such as the rumen or pig gut. ACACT and CoAT are the most divergent loci across the unique RβO operon of *Peptonella*, suggesting their role as molecular determinants of this niche partitioning and highlighting how ecological distinctions may be encoded directly in the core pathway of chain elongators. Finally, differences highlighted by broader comparisons across *Peptonella*, *C. sporogenes*, and *Megasphaera* illustrate how the integration of amino acid catabolism and chain elongation can vary across fermentative lifestyles with meaningful consequences for substrate utilization and product formation.

The integration of amino acid fermentation and chain elongation into a unified metabolic strategy reveals how RβO plays a wider and more ecologically significant role in fermentative physiology than currently recognized. This mechanistic blueprint presents concrete opportunities for strain engineering and bioreactor control to advance MCCA biomanufacturing from protein-rich waste streams. We propose *Peptonella* as a tractable chassis for these efforts, noting that multiple species have already been unknowingly enriched in open-culture waste-to-chemical systems and that the framework described here provides the physiological foundation to deploy them deliberately.

## Materials and Methods

Details on the reagents, catalog numbers, software, and other resource information are provided in supporting information (Table S15).

### Phylogenetics and metabolite data curation

A set of references from the monophyletic portion of Bacillota were identified from the metadata of GTDB^64^ r232, and their genomes were obtained from associated RefSeq annotations (Table S1). Genomes, CDS, and proteins were obtained through NCBI datasets (v18.31.0), totalling 5,953 genomes. A single representative for each genus was retained, numbering 1,108 genomes. Across these representatives, core genes were identified and aligned using GTDBtk^65^ (v2.7.2), and a phylogeny was constructed using IQ-TREE2^66^ with 1000 ultrafast bootstraps, nearest-neighbour interchange optimization (-bnni) and the LG+F+R6 substitution model. The resulting consensus tree was visualized unrooted in iTOL^67^ and labelled with GTDB taxonomy via the internal bulk edit function.

Monocarboxylate production data were retrieved for organisms within Bacillota from metadata reported in a meta-analysis of fermentative prokaryotes^8^. Data from organisms reported to produce straight or branched *n*-butanoate through *n*-octanoate were retained. Experimentally verified instances of MCCA-producing organisms were added to this collection from the literature. Taxonomic information was matched across NCBI-style and GTDB taxonomies, and genus-level representatives were matched to their respective leaves on the consensus tree. Organisms for which confident taxonomic placement could not be obtained, such as when NCBI taxonomy could not be properly resolved against GTDB (e.g., the lack of a whole genome reported for the group), were discarded (supporting information). Representatives for genus-level clades containing known C_4_-C_5_ or ≥C_6_-producing organisms were identified and annotated on the tree via iTOL dataset management. Clades of interest to this work were also manually labelled for visualization purposes.

Smaller trees (e.g., the family-level *Anaerovoracaceae*) were constructed using a similar workflow, but without dereplication to genus-level representatives. These trees were decorated in a similar fashion within iTOL for pathway or gene identities produced via sections below.

### Comparative genomics of MCCA producers

Whole genome sequences from MCCA-producing strains were gathered from reports in the literature, including those recently sequenced from *P. octanoica* and *P. pyruvativorans*^43^. As not all MCCA-producing strains have genomes accessioned in RefSeq, Bakta^68^ (v.1.11.4) was used to predict (pyrodigal [v3.6.3]) and annotate genes uniformly across the collection. Predicted genes were annotated with KofamScan^69^ (v.1.3.0) and eggNOG-mapper^70^ (v.2.1.13; database v.5.0.2). Peptidases were annotated manually by alignment of MCCA producer protein sequences to the MEROPs database using DIAMOND^71^ blastp (--ultra-sensitive) and filtered for best-hits according to bitscore with alignments ≥40% identity and ≥75% alignment overlap retained. Hydrogenases were similarly annotated based on DIAMOND blastp results against the HydDB database^72,73^ (https://github.com/GreeningLab/HydDB) with alignments filtered according to the documentation’s instructions for individual classes of hydrogenases. Pathway analysis was completed with gapseq^74^ (v2.1.0) and the results for enzymes or pathways of interest across MCCA-producing CEB were cross-validated between these results as well as those obtained from KEGG and eggNOG as described above. Resulting annotation information was visualized with MCCA producers sorted and colored by their GTDB-style taxonomic classifications. Additional information on genomic GenBank and RefSeq accessions or annotations results are provided in the supporting information and on the GitHub repository supporting this work.

### Microbial strains

*Peptonella octanoica* F2^43^ served as the model organism for this study. *Peptonella pyruvativorans* (DSM 118842), *Clostridium sporogenes* CE3^43^, *Megasphaera hexanoica* (DSM 106893), *Megasphaera elsdenii* (DSM 20460), and *Caproicibacter fermentans* 7D4C2 (DSM 110548) were also included for comparative experiments.

### Growth media and culture conditions

All strains were cultivated in modified DSM 104 (m104) medium under anaerobic conditions at pH 5-6 and 37 °C. Cultures were incubated either statically or at 150-200 rpm on a shaker. Sodium DL-lactate (TCI or Sigma-Aldrich), casamino acids, glycine, L-alanine, L-proline, sodium acetate, sodium propanoate, and ethanol were added to the medium according to the experimental conditions. Per litre, m104 medium contains 5 g tryptose, 5 g peptone, 10 g yeast extract, 5 g beef extract, 2 g K_2_HPO_4_, 40 mL of salt solution, and 1 mL of 0.1% resazurin. The salt solution (per litre) contains 0.25 g CaCl_2 ·_ 2H_2_O, 0.5 g MgSO_4 ·_ 7H_2_O, 1 g K_2_HPO_4_, 1 g KH_2_PO_4_, 10 g NaHCO_3_, and 2 g NaCl. 10% m104 medium (per litre) contains 0.5 g tryptose, 0.5 g peptone, 1 g yeast extract, 0.5 g beef extract, 2 g K_2_HPO_4_, 40 mL of salt solution, 1 mL of 0.1% resazurin, 1 mL of Wolin’s vitamin solution (10x). The vitamin solution (per litre) contains 20 mg biotin, 20 mg folic acid, 100 mg pyridoxine hydrochloride, 50 mg thiamine HCl, 50 mg riboflavin, 50 mg nicotinic acid, 50 mg calcium D-(+)-pantothenate, 1 mg Vitamin B_12_, 50 mg p-Aminobenzoic acid, and 50 mg (DL)-alpha-Lipoic acid. The vitamin solution was filter-sterilized and stored at 4 °C for later use.

Physiological experiments with *P. octanoica* were conducted in m104 medium at an initial pH of 6 (n = 3 replicates). The following conditions were tested, with supplements added to the medium after autoclaving: (a) no supplements (base medium), (b) 120 mM lactate (TCI), (c) 90 mM proline, (d) 60 mM lactate and 45 mM proline, (e) 60 mM lactate and 30 mM acetate, and (f) 60 mM lactate, 30 mM acetate and 30 mM propanoate. The experiment was conducted in 120 mL serum bottles sealed with rubber stoppers, containing 30 mL of medium and 90 mL of headspace, and incubated in a static incubator.

The utilization of amino acids by *Peptonella* strains and *C. sporogenes* was evaluated using a 10% m104 medium supplemented with 20 g L^-1^ casamino acids in Hungate tubes containing 6 mL of medium and 9 mL of headspace. The initial pH was adjusted to 5, with three replicates conducted. The experimental conditions included, after autoclaving, the addition of: (a) no supplements, (b) 100 mM lactate (TCI), and (c) 100 mM ethanol.

Experiments involving m104 medium supplemented with 100mM glucose, 100 mM acetate, or 100 mM lactate (TCI) to compare microbes used in this study, as well as carbon and amino acid utilization screening for *P. octanoica* strain F2 in 10% m104 medium, were conducted in 96-well plates. The experimental setup used an Agilent BioTek BioSpa live-cell analysis system housed in an anaerobic chamber, coupled to a plate reader for OD600 measurements. Prior to each measurement, plates were subjected to orbital shaking for 1 minute at 2-hour intervals.

The effect of H_2_ on the physiology of *Peptonella* strains was assessed in 10% m104 medium supplemented with 5 g L^-1^ casamino acids and 100 mM sodium DL lactate (Sigma-Aldrich) at an initial pH of 5 in 120 mL serum bottles filled with 45 mL of medium and incubated in a shaking incubator at 200 rpm (n = 4 replicates). Lactate was autoclaved along with the medium. The experimental bottles were (a) pressurized with an H_2_:CO_2_ (80:20) gas mixture to ∼ 1 atm gauge pressure, with pressure released and repressurized every 24 hours to maintain a constant headspace composition throughout, or (b) the headspace pressure was released to ∼ 0 atm gauge pressure every 24 hours.

All synthetic media components (except vitamins) were sterilized by autoclaving at 121 °C for 30 minutes. Strains were inoculated (5% v/v) from glycerol stock into m104 medium containing 50 mM sodium-DL-lactate (25 mM sodium acetate was added to revive *Megasphaera* strains) in Hungate tubes and incubated at 37°C for 3-4 days. The OD600 was normalized across cultures, and a 1% (v/v) inoculum was used for each experimental condition. Samples were collected daily for experiments conducted in serum bottles to measure OD600 and metabolites. Headspace pressure was measured before sampling, and pH was measured at the end of fermentation. OD600 for experiments conducted in Hungate and Balch-type tubes was measured directly with a spectrophotometer (Genesys 30, Thermo Scientific), and samples were collected at the end of each experiment. The samples were stored at –80°C until analysis. Biological replicates across these experiments were assessed for consistency, with one outlier removed for inconsistency in metabolite detection and excluded from further statistical analysis but retained in the supporting information.

Source-separated organics (SSO) or food waste (FW) samples were collected from local full-scale organics processing facilities. The organics were processed as described previously^36^. Briefly, the SSO/FW pulp was sieved through a 0.5 mm sieve, the pH was adjusted to 5-6 using 10N NaOH or 6N HCl, degassed by sparging with N_2_ gas for 1 hour, then transferred to serum bottles, sealed with rubber stoppers, and sterilized by autoclaving. The initial experimental conditions (n = 3 replicates) for testing *P. octanoica* growth were: (a) no supplementation and (b) supplementation with 5 g L^-1^ casamino acids. Later, different batches of SSO with varying concentrations of lactate and SCCAs were tested for *P. octanoica* growth and product profile. The bottles were incubated in a shaking incubator at 150-200 rpm.

Acid whey (AW) was collected from a local dairy processing facility. The pH was adjusted to 5 with 10N NaOH, degassed by sparging with N_2_ for 1 hour, and then pasteurized at 70 °C for 30 minutes. 10 mL of AW was transferred to sterile 25 mL Balch-type tubes within the anaerobic chamber and sealed with a rubber stopper. *P. octanoica* was inoculated into AW and incubated statically with regular headspace pressure monitoring. When headspace pressure accumulated to ∼70 kPa, the pressure was released, and samples were collected to quantify metabolites.

### Lactate, carboxylates and ethanol quantification

Lactate, formate, acetate, and ethanol concentrations were quantified using a high-performance liquid chromatograph (UltiMate 3000 HPLC system, Thermo Scientific, and Agilent 1260 Infinity III HPLC system) with an Aminex HPX-87H column (Bio-Rad, USA). Samples were centrifuged at 10,000g for 10 minutes, diluted with HPLC-grade water, and filtered through a 0.22-μm filter before analysis. The mobile phase was 5 mM sulfuric acid at a constant flow rate of 0.6 mL/min, and 20 μL of sample was injected using the autosampler. The column temperature was set to 50°C, and UV detection was set to 214 nm. Refractive index and UV detection were used to identify and quantify lactate, formate, and acetate. Formate wasn’t detected in any of the *Peptonella* samples. Refractive index detection was used to identify and quantify ethanol.

Alternatively, L– and D-lactate concentrations in sample supernatants were measured using an enzyme-based assay adapted from a previously described method^75^. Briefly, both stereoisomers of lactate were oxidized to pyruvate by their respective lactate dehydrogenases, with NAD⁺ supplied in excess as the electron acceptor. The resulting NADH was quantified spectrophotometrically at 340 nm using a BioTek Epoch 2 microplate spectrophotometer (Agilent), and lactate concentrations were derived from the corresponding NADH signal. The assay was adapted to a 384-well microplate format, with a total reaction volume of 80 µL per well, to increase sample throughput and reduce reagent consumption. All reagents were analytical grade and obtained from Sigma-Aldrich, Bioshop, or Thermo Fisher Scientific.

Carboxylates were quantified using an Agilent 8890 gas chromatograph (GC) equipped with a 7000D Triple Quadrupole mass spectrometer (Agilent), an autosampler (PAL RSI 120 Series 2, Agilent), and a DB-FatWax column (Agilent), with helium as the carrier gas at a flow rate of 1 mL min^-1^. Samples and standards for carboxylic acid analysis were diluted with HPLC-grade water and formic acid (to protonate the carboxylic acids) to a final concentration of 0.1 M, and 0.3 μL was injected at a split ratio of 10:1. The runtime for carboxylate quantification was 16 minutes, with the following temperature gradient: 80°C for 1 minute, a 20°C min^-1^ ramp to 200°C, a 5°C min^-1^ ramp to 210°C, a 20°C min^-1^ ramp to 250°C, then held for 5 minutes. The mass spectrometer was operated in scan mode.

Ethanol was quantified alternately using the same GC-MS system, which operated in headspace mode. Filtered samples were diluted with HPLC-grade water. Before injection, diluted samples and standards were heated at 95 °C for 40 minutes in a tightly sealed vial. The gas phase of the vial was injected into the GC-MS system. The run time was 12 minutes, with the following temperature gradient: 80 °C for 1 minute, a 20 °C min^-1^ ramp to 200 °C, then held for 5 minutes. Helium was used as the carrier gas at a flow rate of 1 mL min^-1^. The mass spectrometer was operated in scan mode.

### Ammonium quantification

Ammonium concentrations were measured using an ion chromatograph (Dionex Integrion HPIC, Thermo Scientific) equipped with a Dionex IonPac CS19 column (Thermo Scientific) and an AS-DV Autosampler. The sample supernatant was diluted using deionized water. 500 μL of diluted samples and standards were transferred into 0.5 mL AS-DV PolyVials (Thermo Scientific) and sealed with filter caps (Thermo Scientific). The mobile phase was deionized water at a constant flow rate of 1 mL min^-1^. The results were analyzed using the Chromeleon software.

### Free amino acid quantification

Free amino acids in the media were derivatized and quantified using the previously described method^76^ with few modifications. The supernatant was filtered using a 0.22 μm filter and diluted with HPLC-grade water. 100 μL of diluted sample was added to a 1.5 mL microfuge tube, and the liquid was evaporated in a Vacufuge Plus (Eppendorf) set to 60 °C for 80 minutes. 70 µL of pyridine and 100 µL of MTBSTFA + 1% TBDMSCl (N-tert-Butyldimethylsilyl-N-methyltrifluoroacetamide with 1% tert-Butyldimethylchlorosilane) were added to dried samples inside a fume hood and mixed by pipetting. The samples were then incubated at 60 °C for 1 hour. The samples were then centrifuged at 14,000g for 1 minute at room temperature, and 150 µL of each supernatant was transferred to a GC-MS vial. Amino acids were quantified using an Agilent 8890 gas chromatograph (GC) equipped with a 7000D Triple Quadrupole mass spectrometer (Agilent), an autosampler (PAL RSI 120 Series 2, Agilent), and a DB-5ms column (Agilent), with helium as the carrier gas at a flow rate of 1 mL min^-1^. 0.5 μL of sample was injected at a split ratio of 10:1. The runtime for amino acid quantification was 45.14 minutes, with the following temperature gradient: 80°C for 2 minutes, a 20°C min^-1^ ramp to 160°C, held for 2 minutes, a 7°C min^-1^ ramp to 280°C, then held for 20 minutes. The mass spectrometer was operated in scan mode.

### Total amino acid quantification with hydrolysis

The total amino acid utilization analysis of *P. octanoica* in m104 medium supplemented with proline was conducted at the SPARC Biocentre, Hospital for Sick Children (SickKids), Toronto, Canada. Total amino acids were determined using the Waters Pico-Tag system, which involves hydrolysis of the dried sample and pre-column derivatization of the hydrolysates with phenylisothiocyanate (PITC) to generate phenylthiocarbamyl (PTC)-amino acids, followed by reverse-phase HPLC analysis. The data were provided to us for further analysis.

The detailed procedure they followed for the analysis is as follows: 1 μg of the supernatant sample was transferred to a 6 x 55 mm borosilicate glass shell vial and dried in a vacuum centrifuge. The dried vials were then placed in a hydrolysis flask containing 300 mL of 6N HCl with 1% phenol. Hydrolysis was carried out at 110 °C for 24 hours under nitrogen. After 24 hours, the shell vials were removed and dried again using a vacuum centrifuge. After drying, samples were treated with a redrying solution consisting of methanol:water:triethylamine (2:2:1), vortex-mixed, and then dried under vacuum (15-20 min). The sample was then derivatized for 15 minutes at room temperature with a derivatizing solution consisting of methanol:water:triethylamine:phenylisothiocyanate (PITC) (7:1:1:1). After 15 minutes, the derivatizing solution was removed under vacuum for 15 minutes. The derivatized sample was then washed with the redrying solution, vortex-mixed, and dried under vacuum for 15 minutes. The derivatized sample was dissolved in a given amount of sample diluent (pH 7.40).

Amino acid analysis was performed on a Waters Acquity UPLC system comprising a binary solvent manager, a sample manager, a TUV detector, and a Waters Acquity UPLC BEH C18 column (2.1 × 100 mm). Data were collected, stored, and processed using Waters Empower 3 chromatography software. Sample aliquots were injected onto the column and resolved using a modified PICO-TAG gradient, with the column temperature set to 48 °C. Derivatized amino acids were detected at 254 nm.

### Carbohydrate quantification

Carbohydrate concentration was estimated using the anthrone method. A total of 2 g of anthrone was dissolved in 1 litre of concentrated H_2_SO_4_ in a fume hood (freshly prepared each time). Glucose served as the standard. Subsequently, 1 mL of the diluted sample was added to a clean tube, followed by 5 mL of the anthrone reagent. The tubes were vortexed, capped, and incubated in a water bath at 90 °C for 17 minutes. After cooling to room temperature, absorbance at 620 nm was measured against the blank.

### Sample preparation for proteomics

Proteomic analysis of *P. octanoica* was conducted on cells grown in mDSM104 medium supplemented with 100 mM lactate and 100 mM proline. Each sample was derived from a 10 mL culture (n=5 per medium condition) grown statically in Hungate tubes. Biomass was collected during the mid-exponential phase by centrifugation at 5000 g for 10 minutes at 4 °C, then washed once with Phosphate-Buffered Saline (PBS, pH 7). Cell pellets were resuspended in 1% sodium deoxycholate in 50 mM triethylammonium bicarbonate (TEAB) buffer at pH 8, and an equal volume of B-PER (Bacterial Protein Extraction Reagent, Thermo Fisher) was then added and mixed. Cells were lysed using 100-µm acid-washed glass beads (Sigma) by bead-beating at 2500 rpm (5 cycles, 1 min each, with 30 s on ice). Samples were then frozen at –80°C to facilitate lysis and thawed slowly on ice. Once thawed, samples were vortexed for 1 minute and centrifuged at 14000 rpm for 10 minutes at 4°C.

Protein quantification was performed using the Pierce™ Bradford Protein Assay Kit (Thermo Fisher) and normalized to the lowest concentration. Next, protein was precipitated with trichloroacetic acid (TCA) at a final concentration of 20% and incubated on ice for 10 minutes. Samples were then centrifuged at 14000 rpm for 5 minutes at 4°C, and the protein pellet was washed twice with ice-cold acetone. Protein pellets were resuspended in 6 M urea in 200 mM ammonium bicarbonate (ABC) buffer. Next, dithiothreitol (DTT) in 200 mM ABC buffer was added to a final concentration of 2.3 mM, and samples were incubated at 37°C for 1 hour. Next, iodoacetamide (IAM) in 200 mM ABC was added to a final concentration of 3.75 mM, and samples were incubated at room temperature in the dark for 30 minutes. Prior to tryptic digestion, samples were diluted with 200 mM ABC to reduce the urea concentration below 1 M. Sequencing-grade trypsin (Promega, Madison, WI, USA) at 2 ng per 1 ug of protein was then added, and the samples were incubated overnight at 37°C and 350 rpm.

Peptide purification was performed using OMIX Tips (Agilent). Trifluoroacetic acid (TFA) was added to the samples to a final concentration of 1%. The OMIX C18 Tip was wetted with a 1:1 acetonitrile (ACN)/H2O mixture, then equilibrated with 0.1% TFA. The tip was washed with 0.1% TFA and eluted with 95% ACN/0.1% formic acid. Samples were dried using a SpeedVac, resuspended in 0.1% formic acid, and filtered through a 0.2 um filter prior to injection.

### LC–MS parameters for peptide separation and detection

All peptide samples were analyzed using a one-dimensional nano-flow LC-MS/MS setup. Briefly, 10 uL of each sample was injected into an EASY-nano LC 1000 system coupled online to a Q-Exactive Orbitrap mass spectrometer (Thermo Scientific) via a nano-electrospray ionization (nanoESI) source. Peptides were separated on a custom-packed SilicaTip emitter column (New Objective, Woburn, USA) packed with ReproSil-Pur C18-AQ resin (Dr. Maisch GmbH, Ammerbuch-Entringen, Baden-Württemberg, Germany). The mobile phases consisted of solvent A (0.1% formic acid in H₂O) and solvent B (0.1% formic acid in acetonitrile), and separation was performed at a constant flow rate of 250 nL/min. A linear gradient was applied as follows: 0 to 5 min, increase to 10% B; 5 to 93 min, increase to 40% B; 93 to 95 min, ramp to 95% B; held at 95% B until 105 min; returned to 0% B by 106 min and held until 120 min for column re-equilibration. The mass spectrometer operated in positive ion mode with a spray voltage of 3 kV and a capillary temperature of 275 °C. Full MS scans were acquired from 400 to 2,000 m/z at a resolution of 70,000, with an AGC target of 1e6 and a maximum injection time of 30 ms. For MS/MS acquisition, a Top10 data-dependent method was employed, isolating the top 10 most intense precursor ions within a 0.4 m/z isolation window. MS2 scans were acquired from 200 to 2,000 m/z at a resolution of 17,500, with a maximum injection time of 50 ms and a normalized collision energy of 27.

### Proteomics data processing

Spectrometry data contained in .raw files were processed with fragpipe^77^ (v23.1) to implement MSFragger-DDA+^78^ (v4.3) and IonQuant^79^ (v1.11.11) with reference proteome prepared via philosopher^80^ (v5.1.2) from the RefSeq annotations of *P. octanoica* strain F2. Using five replicates of each media type, the resulting maximum Label-Free Quantification (maxLFQ) intensities were used to calculate log_2_ fold change (L2FC) and statistical significance was assessed with Welch’s T-test with false discovery correction via Benjamini-Hochberg procedure as implemented in SciPy^81^ and statsmodels, respectively. Volcano plot visualizations were prepared from –log_10_ transform of adjusted p-values and L2FC between m104 with proline and m104 with lactate. Full workflow, tool options, and scripts for downstream statistical analysis are available on the GitHub repository supporting this work. Annotation information, maxLFQ values, and resulting statistics for detected proteins can be found in the supporting information and described in the main text.

### Reconstruction of *Peptonella octanoica* amino acid and lactate catabolism

Results from proteomics were merged with the pathway-level annotations of *P. octanoica* as described earlier in comparative genomics by selecting and manually confirming the identity of proteins as needed to complete major catabolic pathways for amino acid fermentation, portions of central metabolism, and reverse β-oxidation. Protein annotations informed by differential proteomics were tied to associated RefSeq non-redundant protein accession numbers for transparency and are presented in the supporting information (Table S8). When building the metabolic model and its visualization, the expression of each enzyme included was based on confident detection in at least one or more of the media conditions examined proteomically.

To ground annotation of an hydroxyacid dehydrogenase (WP_178263020.1) differentially expressed in the presence of lactate as a putative D-lactate dehydrogenase, sequence alignment against known hydroxyacid dehydrogenases including HicDH, D-LDH, L-LDH, and the catalytic subunit of an electron confurcating L-LDH^22^ were performed via blastp against NCBI’s non-redundant protein collection. Additionally, the tertiary structure similarities of the putative D-LDH and those representing several of these classes of dehydrogenase were investigated. Lastly, AlphaFill^82^ was used to predict ligand locations on the putative D-LDH and provide further visualization. Investigation of the thermodynamic feasibility of lactate and amino acid oxidation was accomplished via eQuilibrator^83^ API with the change in Gibbs free energy plotted across physiologically relevant ranges of electron carrier ratios and intracellular metabolites.

### Comparison of genomic loci encoding RβO enzymes

For comparative analysis of RβO across MCCA producers, within the family *Anaerovoracaceae*, and across the genus *Peptonella*, relevant species-level reference genomes were collected from GTDB or via prior tree-building efforts. Genes encoding RβO enzymes were identified by DIAMOND blastp alignment against the RβO enzymes expressed by *P. octanoica* under lactate-fed conditions. Among MCCA producers, gene synteny was evaluated within alignments in search of collinear blocks containing RβO enzymes. Comparison of gene clusters encoding RβO enzymes within the genus Peptonella were visualized using clinker^84^. Intergenic Rex operator sites were identified using MEME Suite^85^ and compared to a consensus motif (TTGTTAANNNNTTAACAA) reported for *Clostridia*^86^. Visualization of BLAST-like identity of genes encoding RβO enzymes across *Anaerovoracaceae* was accomplished using the tree building methods described above for phylum-level analysis but restricted to the reference genomes within this family plus relevant outgroups and without subsampling. Tree annotation with gene similarity, pathway predictions, and other metadata was accomplished with iTOL.

### Stoichiometry, yield estimation, and product balances

Equivalent concentrations of acetyl-CoA (acetyl-CoA equivalents) were determined using the metabolic reconstruction of *P. octanoica*’s amino acid and lactate catabolism. The contribution of amino acids to acetyl-CoA was taken as the difference between extracellular ammonium concentrations and the sum of observed propanoyl-CoA and branched-chain-CoA products. For product formation, the various monocarboxylates produced by *Peptonella* were assigned different coefficients based on the number of acetyl-CoA involved in their condensation and used to transform observed carboxylate concentrations to acetyl-CoA equivalents. These values are reported in the supporting information (Table S16). This procedure is also documented in scripts handling metabolite analysis on the GitHub supporting this work. Finally, stoichiometric yield coefficients were determined using the same catabolic model. For hydrogen gas yield, a balance of electron carriers was estimated from substrate and product data, with hydrogen production imputed by balancing reduced ferredoxins through RNF complex and ferredoxin-dependent hydrogenase. Terminal carboxylate yields relative to lactate consumption were determined according to Peptonella species. Statistical significance of yields across different experiments was determined by Welch’s t-test.

### Metagenomics of isolated *Peptonella* species

Sandpiper^48^ was used to identify short-read metagenomic datasets with detectable *P. octanoica* and *P. pyruvativorans* signal, with each of the SRA accessions flagged by the tool undergoing manual review to determine environmental context of the associated metagenome. To examine population-level differences, datasets were obtained with SRA Toolkit’s fasterq-dump, quality trimmed with fastp^87^ (v1.1.0), mapped to the reference genome *of P. octanoica* F2 via BWA-MEM2^88^ (v2.3), and statistics summarized with CoverM^89^ (v0.7.0). Sample accessions, environmental contexts, and both abundance metrics reported by Sandpiper and this analysis are contained in supporting information (Table S13).

## Data Availability

Proteomic data are available on the PRIDE server (PXD083949). Accession numbers for all genomic and metagenomic data used herein are provided in the supporting information. Scripts for data analysis and a reduced complexity genome-scale metabolic model of *Peptonella* lactate and amino acid metabolism are available on GitHub (https://github.com/bglindner/proteins-to-MCCAs).

## Acknowledgements

We are grateful for Jasmeen Parmar and Diana Dyussekenova’s contributions to food waste collection and acknowledge the City of Toronto for providing sampling access to organics processing facilities. We thank Elizabeth Lee-Chin for assistance with early experiments related to this work, and Robert Flick for running LC-MS analyses and supporting GC-MS operations for amino acid quantification. This work was supported by the Natural Sciences and Engineering Research Council of Canada (RGPIN-2021-02684, NSERC-CREATE 528163-2019). Computational analyses were supported in part by infrastructure maintained by the Digital Research Alliance of Canada.

## Citations

1. Jackson, B.E., and McInerney, M.J. (2002). Anaerobic microbial metabolism can proceed close to thermodynamic limits. Nature 415, 454–456. 10.1038/415454a.

2. Singh, V., Lee, G., Son, H., Koh, H., Kim, E.S., Unno, T., and Shin, J.-H. (2022). Butyrate producers, “The Sentinel of Gut”: Their intestinal significance with and beyond butyrate, and prospective use as microbial therapeutics. Front Microbiol 13, 1103836. 10.3389/fmicb.2022.1103836.

3. Louis, P., and Flint, H.J. (2009). Diversity, metabolism and microbial ecology of butyrate-producing bacteria from the human large intestine. FEMS Microbiol Lett 294, 1–8. 10.1111/j.1574-6968.2009.01514.x.

4. Storm, A.C., Kristensen, N.B., and Hanigan, M.D. (2012). A model of ruminal volatile fatty acid absorption kinetics and rumen epithelial blood flow in lactating Holstein cows. Journal of Dairy Science 95, 2919–2934. 10.3168/jds.2011-4239.

5. Rappé, M.S., and Giovannoni, S.J. (2003). The Uncultured Microbial Majority. Annu. Rev. Microbiol. 57, 369–394. 10.1146/annurev.micro.57.030502.090759.

6. Rinke, C., Schwientek, P., Sczyrba, A., Ivanova, N.N., Anderson, I.J., Cheng, J.-F., Darling, A., Malfatti, S., Swan, B.K., Gies, E.A., et al. (2013). Insights into the phylogeny and coding potential of microbial dark matter. Nature 499, 431–437. 10.1038/nature12352.

7. Nauwynck, W., Sakarika, M., Faust, K., and Boon, N. (2025). Differential recovery of chain-elongating bacteria: comparing droplet, plating, and dilution-to-extinction methods. mSystems 10, e0135625. 10.1128/msystems.01356-25.

8. Hackmann, T.J., and Zhang, B. (2023). The phenotype and genotype of fermentative prokaryotes. Science Advances 9, eadg8687. 10.1126/sciadv.adg8687.

9. Hackmann, T.J. (2024). The vast landscape of carbohydrate fermentation in prokaryotes. FEMS Microbiol Rev 48, fuae016. 10.1093/femsre/fuae016.

10. Candry, P., and Ganigué, R. (2021). Chain elongators, friends, and foes. Current Opinion in Biotechnology 67, 99–110. 10.1016/j.copbio.2021.01.005.

11. Sola, L., Candeliere, F., Busi, E., Raimondi, S., Amaretti, A., and Rossi, M. (2026). A genomic atlas of gut clostridia: phylogeny, butyrate, and propionate production. Front. Microbiol. 17. 10.3389/fmicb.2026.1761627.

12. Tahmid, M., Choi, H.J., Cruz, A., Ferreira-Garcia, D., Abbasi, E., Botte, G., and Hatzell, M. Waste-Based Volatile Fatty Acids for Fuel and Chemical Production. ChemRxiv 2026. 10.26434/chemrxiv-2026-55d4t.

13. Agena, E., Gois, I.M., Bowers, C.M., Mahadevan, R., Scarborough, M.J., and Lawson, C.E. (2024). Evaluating the feasibility of medium-chain oleochemical synthesis using microbial chain elongation. J. Ind. Microbiol. Biotechnol. 51, kuae027. 10.1093/jimb/kuae027.

14. Barker, H.A. (1937). The production of caproic and butyric acids by the methane fermentation of ethyl alcohol. Archiv. Mikrobiol. 8, 415–421. 10.1007/BF00407210.

15. Barker, H.A., and Taha, S.M. (1942). Clostridium kluyverii, an Organism Concerned in the Formation of Caproic Acid from Ethyl Alcohol. Journal of Bacteriology 43, 347–363. 10.1128/jb.43.3.347-363.1942.

16. Bornstein, B.T., and Barker, H.A. (1948). THE ENERGY METABOLISM OF CLOSTRIDIUM KLUYVERI AND THE SYNTHESIS OF FATTY ACIDS. Journal of Biological Chemistry 172, 659–669. 10.1016/S0021-9258(19)52752-1.

17. Seedorf, H., Fricke, W.F., Veith, B., Brüggemann, H., Liesegang, H., Strittmatter, A., Miethke, M., Buckel, W., Hinderberger, J., Li, F., et al. (2008). The genome of Clostridium kluyveri, a strict anaerobe with unique metabolic features. Proceedings of the National Academy of Sciences 105, 2128–2133. 10.1073/pnas.0711093105.

18. Thauer, R.K., Jungermann, K., and Decker, K. (1977). Energy conservation in chemotrophic anaerobic bacteria. Bacteriological Reviews 41, 100–180. 10.1128/br.41.1.100-180.1977.

19. Angenent, L.T., Richter, H., Buckel, W., Spirito, C.M., Steinbusch, K.J.J., Plugge, C.M., Strik, D.P.B.T.B., Grootscholten, T.I.M., Buisman, C.J.N., and Hamelers, H.V.M. (2016). Chain Elongation with Reactor Microbiomes: Open-Culture Biotechnology To Produce Biochemicals. Environ. Sci. Technol. 50, 2796–2810. 10.1021/acs.est.5b04847.

20. Xu, J., Hao, J., Guzman, J.J.L., Spirito, C.M., Harroff, L.A., and Angenent, L.T. (2018). Temperature-Phased Conversion of Acid Whey Waste Into Medium-Chain Carboxylic Acids via Lactic Acid: No External e-Donor. Joule 2, 280–295. 10.1016/j.joule.2017.11.008.

21. Contreras-Dávila, C.A., Carrión, V.J., Vonk, V.R., Buisman, C.N.J., and Strik, D.P.B.T.B. (2020). Consecutive lactate formation and chain elongation to reduce exogenous chemicals input in repeated-batch food waste fermentation. Water Research 169, 115215. 10.1016/j.watres.2019.115215.

22. Weghoff, M.C., Bertsch, J., and Müller, V. (2015). A novel mode of lactate metabolism in strictly anaerobic bacteria. Environmental Microbiology 17, 670–677. 10.1111/1462-2920.12493.

23. Schink, B. (2015). Electron confurcation in anaerobic lactate oxidation. Environmental Microbiology 17, 543–543. 10.1111/1462-2920.12568.

24. Gois, I.M., Bowers, C.M., Kim, B.-C., Flick, R., and Lawson, C.E. (2026). Acetate utilization strategy in chain-elongating bacteria determines butyrate versus medium-chain carboxylate production. Nat Microbiol, 1–13. 10.1038/s41564-026-02320-8.

25. Jeon, B.S., Kim, E.-J., Seo, H., Kim, H., Shin, S., Schlaiß, C., Angenent, L.T., Kim, K.-J., and Sang, B.-I. (2025). Molecular Chain Elongation Mechanism for n-Caproate Biosynthesis by Megasphaera Hexanoica. Advanced Science 12, e06069. 10.1002/advs.202506069.

26. Sakarika, M., Regueira, A., Benson, D., Singh, A., and Ganigué, R. (2026). The choice of electron donor determines the product spectrum in thermophilic chain elongation. Journal of Environmental Chemical Engineering, 124605. 10.1016/j.jece.2026.124605.

27. Cabral, L. da S., and Weimer, P.J. (2024). Megasphaera elsdenii: Its Role in Ruminant Nutrition and Its Potential Industrial Application for Organic Acid Biosynthesis. Microorganisms 12, 219. 10.3390/microorganisms12010219.

28. Wallace, R.J. (1986). Catabolism of Amino Acids by Megasphaera elsdenii LC1. Applied and Environmental Microbiology 51, 1141–1143. 10.1128/aem.51.5.1141-1143.1986.

29. Rychlik, J.L., LaVera, R., and Russell, J.B. (2002). Amino Acid Deamination by Ruminal Megasphaera elsdenii Strains. Curr Microbiol 45, 340–345. 10.1007/s00284-002-3743-4.

30. Wallace, R.J., McKain, N., McEwan, N.R., Miyagawa, E., Chaudhary, L.C., King, T.P., Walker, N.D., Apajalahti, J.H.A., and Newbold, C.J. (2003). Eubacterium pyruvativorans sp. nov., a novel non-saccharolytic anaerobe from the rumen that ferments pyruvate and amino acids, forms caproate and utilizes acetate and propionate. Int J Syst Evol Microbiol 53, 965– 970. 10.1099/ijs.0.02110-0.

31. Wallace, R.J., Chaudhary, L.C., Miyagawa, E., McKain, N., and Walker, N.D. (2004). Metabolic properties of Eubacterium pyruvativorans, a ruminal ‘hyper-ammonia-producing’ anaerobe with metabolic properties analogous to those of Clostridium kluyveri. Microbiology 150, 2921–2930. 10.1099/mic.0.27190-0.

32. Strachan, C.R., Bowers, C.M., Kim, B.-C., Movsesijan, T., Neubauer, V., Mueller, A.J., Yu, X.A., Pereira, F.C., Nagl, V., Faas, J., et al. (2025). Distinct lactate utilization strategies drive niche differentiation between two co-existing Megasphaera species in the rumen microbiome. ISME J 19, wraf147. 10.1093/ismejo/wraf147.

33. Joshi, S., Robles, A., Aguiar, S., and Delgado, A.G. (2021). The occurrence and ecology of microbial chain elongation of carboxylates in soils. ISME J 15, 1907–1918. 10.1038/s41396-021-00893-2.

34. Wang, H., Jeon, B.S., Ortiz-Ardila, A.E., and Angenent, L.T. (2025). Steering the Chain-Elongating Microbiome to C8 or C6 Production with Ethanol and Lactate as Coelectron Donors. Environ. Sci. Technol. 59, 16947–16958. 10.1021/acs.est.5c01461.

35. Candry, P., Flinkstrom, Z., and Henriikka Winkler, M.-K. Wetlands harbor lactic acid-driven chain elongators. Microbiol Spectr 12, e02105–23. 10.1128/spectrum.02105-23.

36. Dyussekenova, D., Parmar, J.K., Ezabadi, M.A., Lindner, B.G., Hong, Y., Werber, J.R., and Lawson, C.E. (2026). Long-Term Production and Recovery of Medium-Chain Carboxylates from Source-Separated Organics. ACS EST Eng. 10.1021/acsestengg.6c00464.

37. Albright, S., and Louca, S. (2023). Trait biases in microbial reference genomes. Sci Data 10, 84. 10.1038/s41597-023-01994-7.

38. Vital, M., Howe, A.C., and Tiedje, J.M. (2014). Revealing the Bacterial Butyrate Synthesis Pathways by Analyzing (Meta)genomic Data. mBio 5, 10.1128/mbio.00889-14. https://doi.org/10.1128/mbio.00889-14.

39. Neumann-Schaal, M., Jahn, D., and Schmidt-Hohagen, K. (2019). Metabolism the Difficile Way: The Key to the Success of the Pathogen Clostridioides difficile. Front. Microbiol. 10. 10.3389/fmicb.2019.00219.

40. Liu, Y., Chen, H., Van Treuren, W., Hou, B.-H., Higginbottom, S.K., and Dodd, D. (2022). Clostridium sporogenes uses reductive Stickland metabolism in the gut to generate ATP and produce circulating metabolites. Nat Microbiol 7, 695–706. 10.1038/s41564-022-01109-9.

41. Fonknechten, N., Chaussonnerie, S., Tricot, S., Lajus, A., Andreesen, J.R., Perchat, N., Pelletier, E., Gouyvenoux, M., Barbe, V., Salanoubat, M., et al. (2010). Clostridium sticklandii, a specialist in amino acid degradation:revisiting its metabolism through its genome sequence. BMC Genomics 11, 555. 10.1186/1471-2164-11-555.

42. Pavao, A., Graham, M., Arrieta-Ortiz, M., Immanuel, S.R.C., Baliga, N.S., and Bry, L. (2022). Reconsidering the in vivo functions of Clostridial Stickland amino acid fermentations. Anaerobe 76, 102600. 10.1016/j.anaerobe.2022.102600.

43. Nallasamy, D.K., Lindner, B.G., and Lawson, C.E. (2026). Peptonella octanoica gen. nov., sp. nov., a new medium-chain carboxylate-producing bacterium, and the reclassification of Eubacterium pyruvativorans as Peptonella pyruvativorans comb. nov. Preprint at bioRxiv, 10.64898/2026.08.23.746564 https://doi.org/10.64898/2026.08.23.746564.

44. Kayastha, K., Katsyv, A., Himmrich, C., Welsch, S., Schuller, J.M., Ermler, U., and Müller, V. (2022). Structure-based electron-confurcation mechanism of the Ldh-EtfAB complex. eLife 11, e77095. 10.7554/eLife.77095.

45. Guo, C.-J., Allen, B.M., Hiam, K.J., Dodd, D., Van Treuren, W., Higginbottom, S., Nagashima, K., Fischer, C.R., Sonnenburg, J.L., Spitzer, M.H., et al. (2019). Depletion of microbiome-derived molecules in the host using Clostridium genetics. Science 366, eaav1282. 10.1126/science.aav1282.

46. Louis, P., McCrae, S.I., Charrier, C., and Flint, H.J. (2007). Organization of butyrate synthetic genes in human colonic bacteria: phylogenetic conservation and horizontal gene transfer. FEMS Microbiol Lett 269, 240–247. 10.1111/j.1574-6968.2006.00629.x.

47. Rabinowitz, J.D., and Enerbäck, S. (2020). Lactate: the ugly duckling of energy metabolism. Nat Metab 2, 566–571. 10.1038/s42255-020-0243-4.

48. Woodcroft, B.J., Aroney, S.T.N., Zhao, R., Cunningham, M., Mitchell, J.A.M., Nurdiansyah, R., Blackall, L., and Tyson, G.W. (2026). Comprehensive taxonomic identification of microbial species in metagenomic data using SingleM and Sandpiper. Nat Biotechnol 44, 948–953. 10.1038/s41587-025-02738-1.

49. United Nations Environment Programme Food Waste Index Report 2024.

50. Gatto, A., and Chepeliev, M. (2024). Global food loss and waste estimates show increasing nutritional and environmental pressures. Nat Food 5, 136–147. 10.1038/s43016-023-00915-6.

51. Zhu, J., Luo, Z., Sun, T., Li, W., Zhou, W., Wang, X., Fei, X., Tong, H., and Yin, K. (2023). Cradle-to-grave emissions from food loss and waste represent half of total greenhouse gas emissions from food systems. Nat Food 4, 247–256. 10.1038/s43016-023-00710-3.

52. Roy, P., Mohanty, A.K., Dick, P., and Misra, M. (2023). A Review on the Challenges and Choices for Food Waste Valorization: Environmental and Economic Impacts. ACS Environ. Au 3, 58–75. 10.1021/acsenvironau.2c00050.

53. Pavao, A., Graham, M., Arrieta-Ortiz, M., Immanuel, S.R.C., Baliga, N.S., and Bry, L. (2022). Reconsidering the in vivo functions of Clostridial Stickland amino acid fermentations. Anaerobe 76, 102600. 10.1016/j.anaerobe.2022.102600.

54. Lopez, C.A., McNeely, T.P., Nurmakova, K., Beavers, W.N., and Skaar, E.P. (2020). *Clostridioides difficile* proline fermentation in response to commensal clostridia. Anaerobe 63, 102210. 10.1016/j.anaerobe.2020.102210.

55. Moore, A.E., Leclercq, G.M.E., Moffatt, L.S., Fitzgerald, B.G., and Sorbara, M.T. (2026). Differential use of Stickland fermentation by *Lachnospiraceae* regulates *Clostridioides difficile* growth. Gut Microbiology 3, 100015. 10.1016/j.gutmic.2026.100015.

56. Putumbaka, S., Schut, G.J., Thorgersen, M.P., Poole, F.L., Shao, N., Rodionov, D.A., and Adams, M.W.W. (2025). Tungsten is utilized for lactate consumption and SCFA production by a dominant human gut microbe Eubacterium limosum. Proceedings of the National Academy of Sciences 122, e2411809121. 10.1073/pnas.2411809121.

57. Köpke, M., Straub, M., and Dürre, P. (2013). Clostridium difficile is an autotrophic bacterial pathogen. PLoS One 8, e62157. 10.1371/journal.pone.0062157.

58. Zheng, Y., Kahnt, J., Kwon, I.H., Mackie, R.I., and Thauer, R.K. (2014). Hydrogen Formation and Its Regulation in Ruminococcus albus: Involvement of an Electron-Bifurcating [FeFe]-Hydrogenase, of a Non-Electron-Bifurcating [FeFe]-Hydrogenase, and of a Putative Hydrogen-Sensing [FeFe]-Hydrogenase. Journal of Bacteriology 196, 3840–3852. 10.1128/jb.02070-14.

59. Andreesen, J.R. (1994). Glycine metabolism in anaerobes. Antonie van Leeuwenhoek 66, 223–237. 10.1007/BF00871641.

60. Bouillaut, L., Dubois, T., Francis, M.B., Daou, N., Monot, M., Sorg, J.A., Sonenshein, A.L., and Dupuy, B. (2019). Role of the global regulator Rex in control of NAD+-regeneration in Clostridioides (Clostridium) difficile. Molecular Microbiology 111, 1671– 1688. 10.1111/mmi.14245.

61. Bouillaut, L., Self, W.T., and Sonenshein, A.L. (2013). Proline-Dependent Regulation of Clostridium difficile Stickland Metabolism. J Bacteriol 195, 844–854. 10.1128/JB.01492-12.

62. Baleeiro, F.C.F., Kleinsteuber, S., and Sträuber, H. (2021). Hydrogen as a Co-electron Donor for Chain Elongation With Complex Communities. Front. Bioeng. Biotechnol. 9. 10.3389/fbioe.2021.650631.

63. Olson, S.T., and Massey, V. (1979). Purification and properties of the flavoenzyme D-lactate dehydrogenase from Megasphaera elsdenii. Biochemistry 18, 4714–4724. 10.1021/bi00588a036.

64. Parks, D.H., Chaumeil, P.-A., Mussig, A.J., Rinke, C., Chuvochina, M., and Hugenholtz, P. (2026). GTDB release 10: a complete and systematic taxonomy for 715 230 bacterial and 17 245 archaeal genomes. Nucleic Acids Res 54, D743–D754. 10.1093/nar/gkaf1040.

65. Chaumeil, P.-A., Mussig, A.J., Hugenholtz, P., and Parks, D.H. (2020). GTDB-Tk: a toolkit to classify genomes with the Genome Taxonomy Database. Bioinformatics 36, 1925– 1927. 10.1093/bioinformatics/btz848.

66. Nguyen, L.-T., Schmidt, H.A., von Haeseler, A., and Minh, B.Q. (2015). IQ-TREE: A Fast and Effective Stochastic Algorithm for Estimating Maximum-Likelihood Phylogenies. Mol Biol Evol 32, 268–274. 10.1093/molbev/msu300.

67. Letunic, I., and Bork, P. (2021). Interactive Tree Of Life (iTOL) v5: an online tool for phylogenetic tree display and annotation. Nucleic Acids Res 49, W293–W296. 10.1093/nar/gkab301.

68. Schwengers, O., Jelonek, L., Dieckmann, M.A., Beyvers, S., Blom, J., and Goesmann, A. (2021). Bakta: rapid and standardized annotation of bacterial genomes via alignment-free sequence identification. Microbial Genomics 7, 000685. 10.1099/mgen.0.000685.

69. Aramaki, T., Blanc-Mathieu, R., Endo, H., Ohkubo, K., Kanehisa, M., Goto, S., and Ogata, H. (2020). KofamKOALA: KEGG Ortholog assignment based on profile HMM and adaptive score threshold. Bioinformatics 36, 2251–2252. 10.1093/bioinformatics/btz859.

70. Cantalapiedra, C.P., Hernández-Plaza, A., Letunic, I., Bork, P., and Huerta-Cepas, J. (2021). eggNOG-mapper v2: Functional Annotation, Orthology Assignments, and Domain Prediction at the Metagenomic Scale. Mol Biol Evol 38, 5825–5829. 10.1093/molbev/msab293.

71. Buchfink, B., Xie, C., and Huson, D.H. (2015). Fast and sensitive protein alignment using DIAMOND. Nat Methods 12, 59–60. 10.1038/nmeth.3176.

72. Søndergaard, D., Pedersen, C.N.S., and Greening, C. (2016). HydDB: A web tool for hydrogenase classification and analysis. Sci Rep 6, 34212. 10.1038/srep34212.

73. Greening, C., Biswas, A., Carere, C.R., Jackson, C.J., Taylor, M.C., Stott, M.B., Cook, G.M., and Morales, S.E. (2016). Genomic and metagenomic surveys of hydrogenase distribution indicate H2 is a widely utilised energy source for microbial growth and survival. ISME J 10, 761–777. 10.1038/ismej.2015.153.

74. Zimmermann, J., Kaleta, C., and Waschina, S. (2021). gapseq: informed prediction of bacterial metabolic pathways and reconstruction of accurate metabolic models. Genome Biol 22, 81. 10.1186/s13059-021-02295-1.

75. Hass, D., and Hurley, J. (2023). Lactate Concentration assay (LDH method).

76. Long, C.P., and Antoniewicz, M.R. (2019). High-resolution 13C metabolic flux analysis. Nat Protoc 14, 2856–2877. 10.1038/s41596-019-0204-0.

77. Hsiao, Y., Zhang, H., Li, G.X., Deng, Y., Yu, F., Valipour Kahrood, H., Steele, J.R., Schittenhelm, R.B., and Nesvizhskii, A.I. (2024). Analysis and Visualization of Quantitative Proteomics Data Using FragPipe-Analyst. J. Proteome Res. 23, 4303–4315. 10.1021/acs.jproteome.4c00294.

78. Yu, F., Deng, Y., and Nesvizhskii, A.I. (2025). MSFragger-DDA+ enhances peptide identification sensitivity with full isolation window search. Nat Commun 16, 3329. 10.1038/s41467-025-58728-z.

79. Yu, F., Haynes, S.E., and Nesvizhskii, A.I. (2021). IonQuant Enables Accurate and Sensitive Label-Free Quantification With FDR-Controlled Match-Between-Runs. Molecular & Cellular Proteomics 20. 10.1016/j.mcpro.2021.100077.

80. da Veiga Leprevost, F., Haynes, S.E., Avtonomov, D.M., Chang, H.-Y., Shanmugam, A.K., Mellacheruvu, D., Kong, A.T., and Nesvizhskii, A.I. (2020). Philosopher: a versatile toolkit for shotgun proteomics data analysis. Nat Methods 17, 869–870. 10.1038/s41592-020-0912-y.

81. Virtanen, P., Gommers, R., Oliphant, T.E., Haberland, M., Reddy, T., Cournapeau, D., Burovski, E., Peterson, P., Weckesser, W., Bright, J., et al. (2020). SciPy 1.0: fundamental algorithms for scientific computing in Python. Nat Methods 17, 261–272. 10.1038/s41592-019-0686-2.

82. Hekkelman, M.L., de Vries, I., Joosten, R.P., and Perrakis, A. (2023). AlphaFill: enriching AlphaFold models with ligands and cofactors. Nat Methods 20, 205–213. 10.1038/s41592-022-01685-y.

83. Beber, M.E., Gollub, M.G., Mozaffari, D., Shebek, K.M., Flamholz, A.I., Milo, R., and Noor, E. (2021). eQuilibrator 3.0: a database solution for thermodynamic constant estimation. Nucleic Acids Res 50, D603–D609. 10.1093/nar/gkab1106.

84. Gilchrist, C.L.M., and Chooi, Y.-H. (2021). clinker & clustermap.js: automatic generation of gene cluster comparison figures. Bioinformatics 37, 2473–2475. 10.1093/bioinformatics/btab007.

85. Bailey, T.L., Boden, M., Buske, F.A., Frith, M., Grant, C.E., Clementi, L., Ren, J., Li, W.W., and Noble, W.S. (2009). MEME SUITE: tools for motif discovery and searching. Nucleic Acids Res 37, W202–208. 10.1093/nar/gkp335.

86. Zhang, L., Nie, X., Ravcheev, D.A., Rodionov, D.A., Sheng, J., Gu, Y., Yang, S., Jiang, W., and Yang, C. (2014). Redox-Responsive Repressor Rex Modulates Alcohol Production and Oxidative Stress Tolerance in Clostridium acetobutylicum. J Bacteriol 196, 3949–3963. 10.1128/JB.02037-14.

87. Chen, S., Zhou, Y., Chen, Y., and Gu, J. (2018). fastp: an ultra-fast all-in-one FASTQ preprocessor. Bioinformatics 34, i884–i890. 10.1093/bioinformatics/bty560.

88. Vasimuddin, Md., Misra, S., Li, H., and Aluru, S. (2019). Efficient Architecture-Aware Acceleration of BWA-MEM for Multicore Systems. In 2019 IEEE International Parallel and Distributed Processing Symposium (IPDPS), pp. 314–324. 10.1109/IPDPS.2019.00041.

89. Aroney, S.T.N., Newell, R.J.P., Nissen, J.N., Camargo, A.P., Tyson, G.W., and Woodcroft, B.J. (2025). CoverM: read alignment statistics for metagenomics. Bioinformatics 41, btaf147. 10.1093/bioinformatics/btaf147.

