## Supplementary material for "Coupling Stickland fermentation with reverse β-oxidation represents a new mode of medium-chain carboxylate production": Supp Data 1

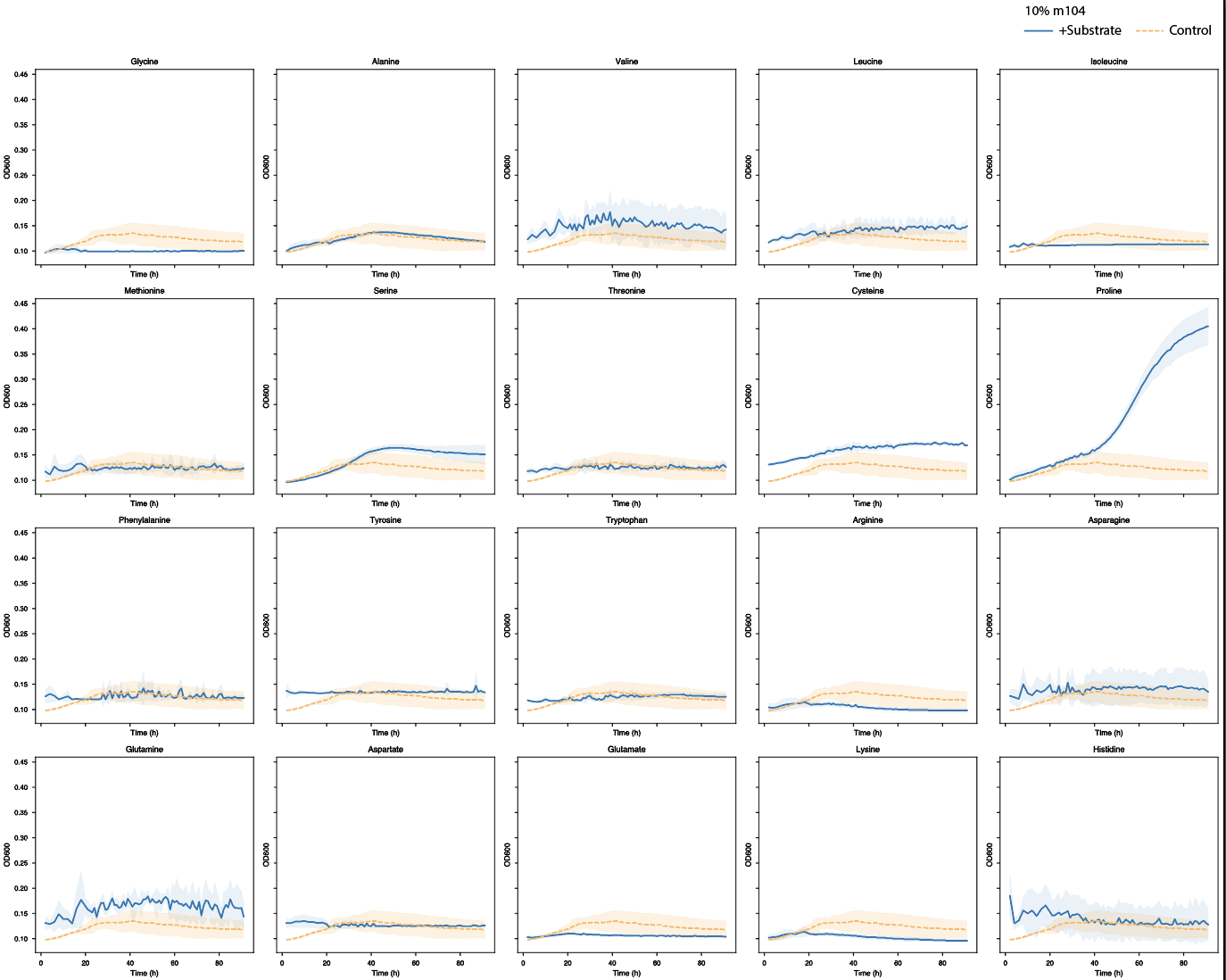

**Figure S1**. High throughput substrate utilization screening results in light formulation of m104. Growth curves were generated with three replicates across 20 substrates. Control growth curve represents 10% m104 without any additions.

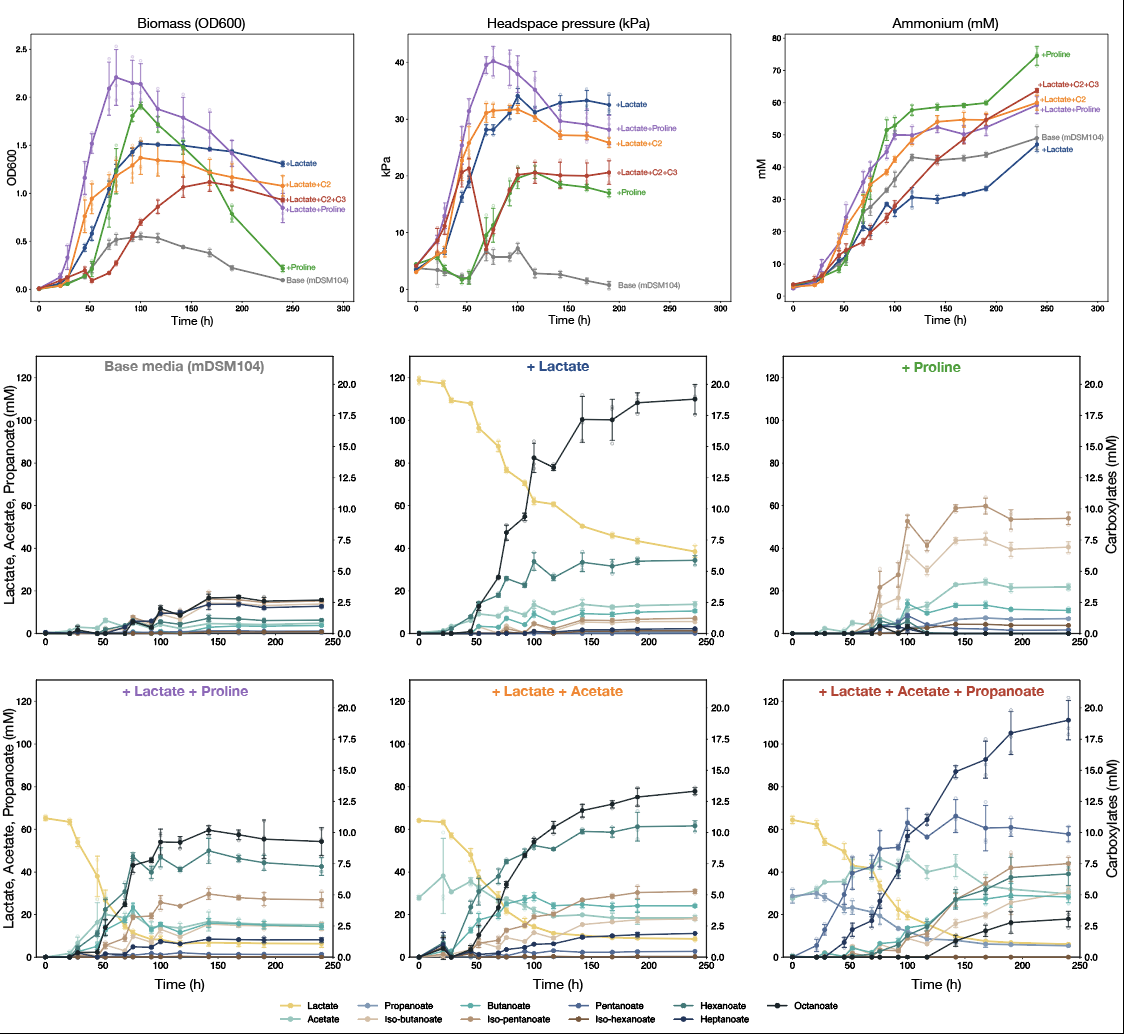

**Figure S2.** The influence of various substrates on *P. octanoica’*s growth and metabolite formation in m104. Growth, headspace pressure, and deamination curves (top row) of *P. octanoica* grown across six formulations of m104 (indicated with labels). Metabolite formation across each media composition (middle and bottom row). Points are the averages of biological replicates (n=3) and error bars represent one standard deviation.

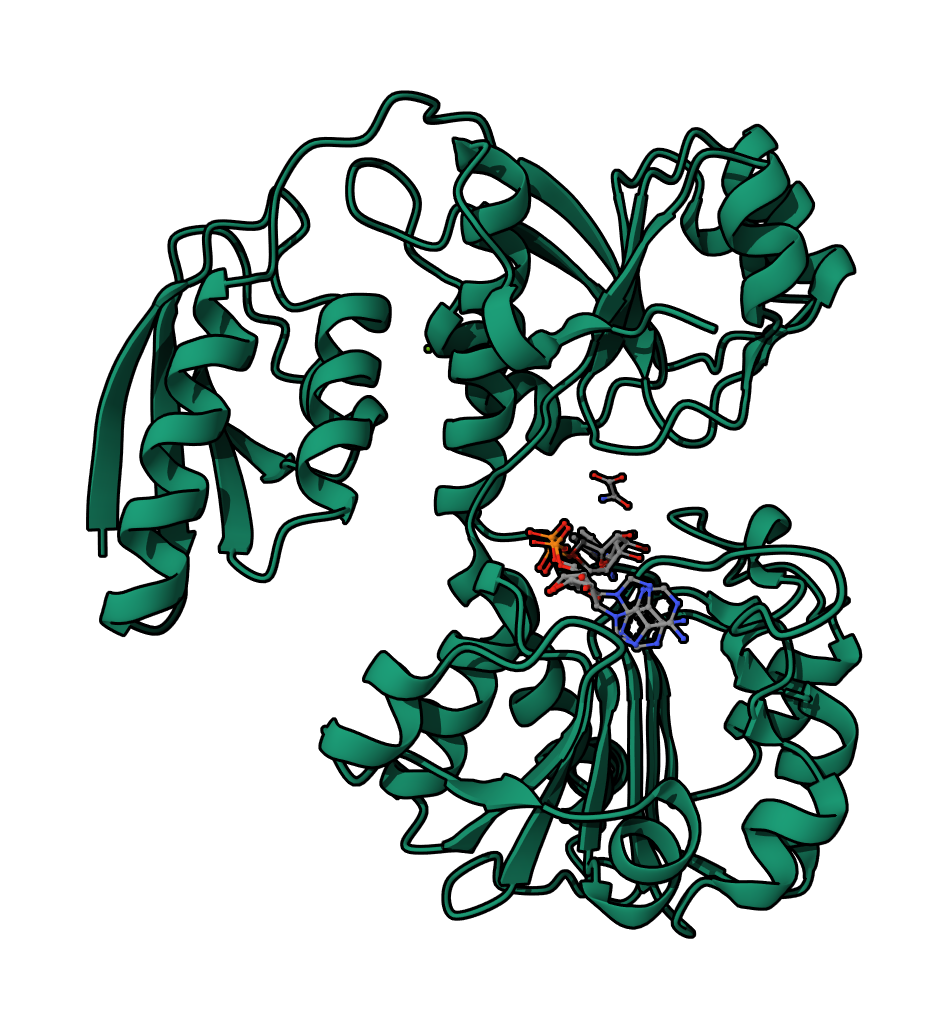

**Figure S3.** A putative D-LDH with a fused HPr domain identified with proteomics and differentially expressed under lactate-fed conditions. The structure for this enzyme is shown with AlphaFill predicted substrates including oxamate (a pyruvate proxy) adjacent to NAD^+^ / NADH.

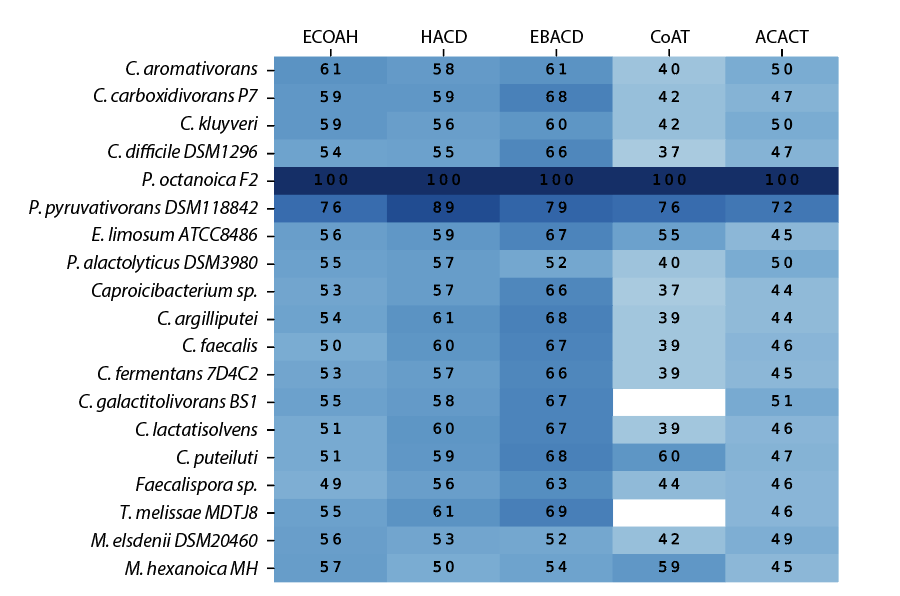

**Figure S4.** Homology across RBO-encoding genes across MCCA producers using proteomically identified references from P*. octanoica*. Shading and printed values in the heatmap represent amino acid identities.

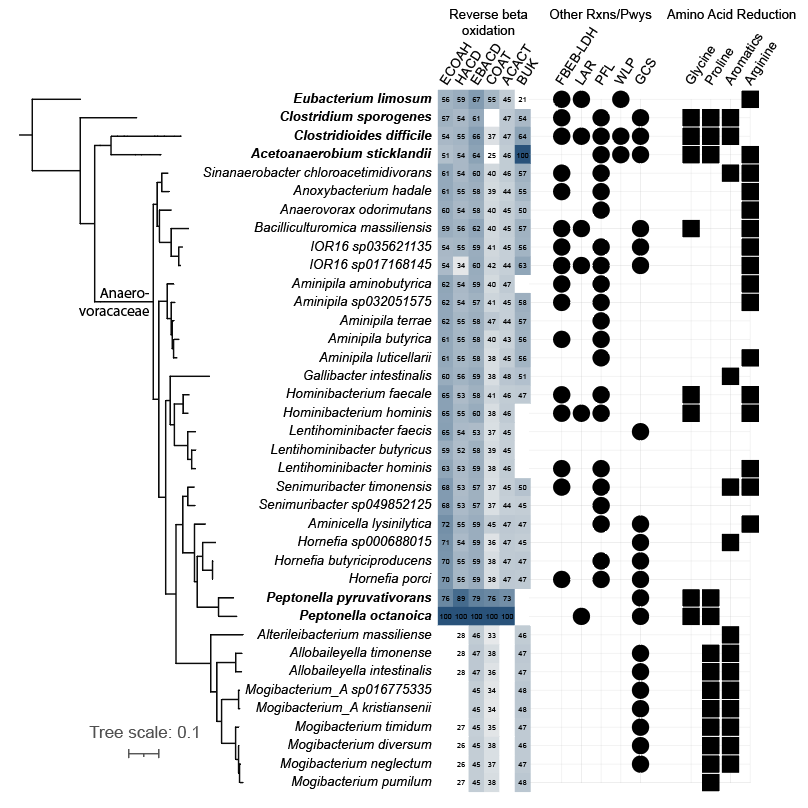

**Figure S5.** Stickland pathways and RBO enzymes are broadly distributed across the family *Anaerovoracaceae*.

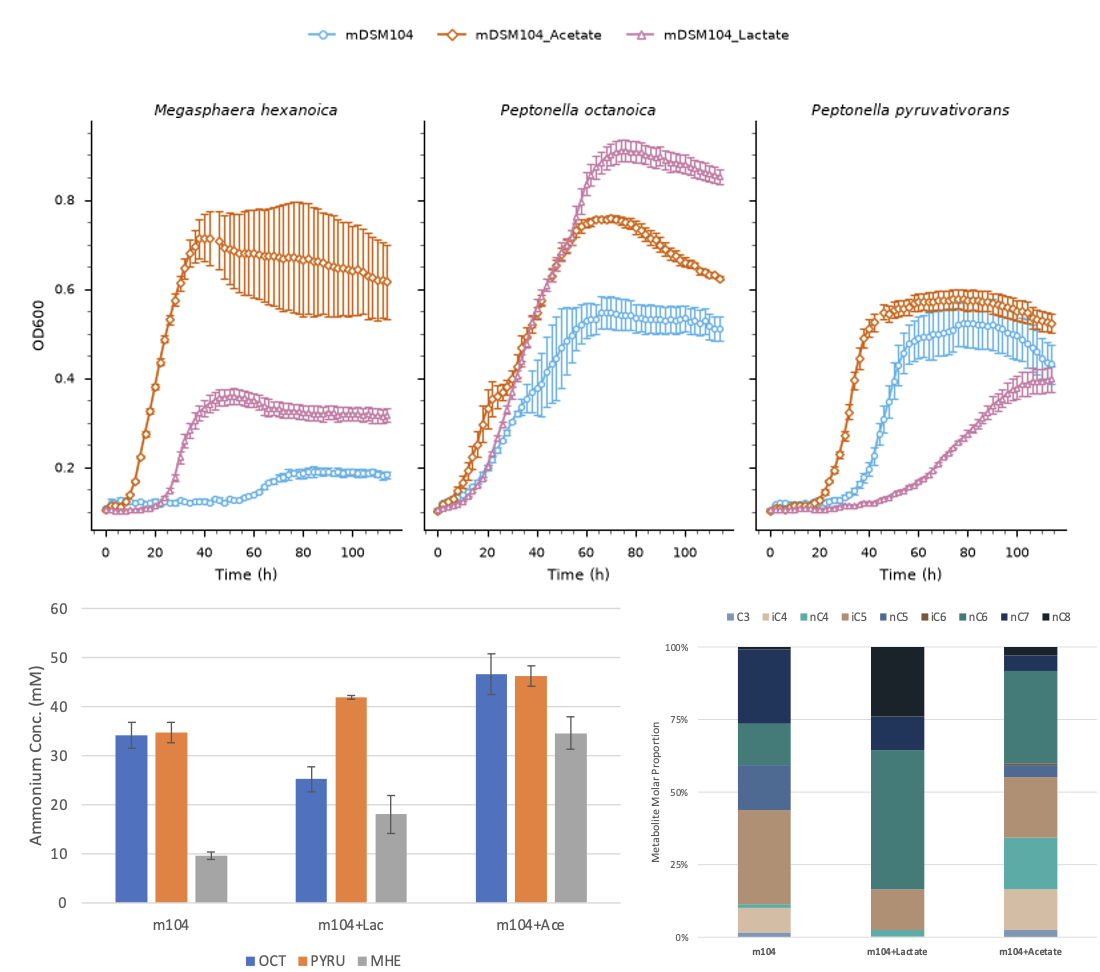

**Figure S6**. Comparative growth and metabolite formation of *M. hexanoica* and *Peptonella*. Three media compositions were used: basal m104, m104 with acetate, and m104 with L-lactate. Growth curves (top panel) for the three species are coloured according to media types. Points are the average of biological replicates (n=3) and error bars represent one standard deviation. Endpoint ammonium concentrations (bottom left) for each species are also reported as averages and standard deviations. Average product carboxylate profiles (bottom right) are shown for *M. hexanoica* scaled to molar proportions across the three media types. Abbreviations include OCT (*P. octanoica*), PYRU (*P. pyruvativorans*), and MHE (*M. hexanoica*).

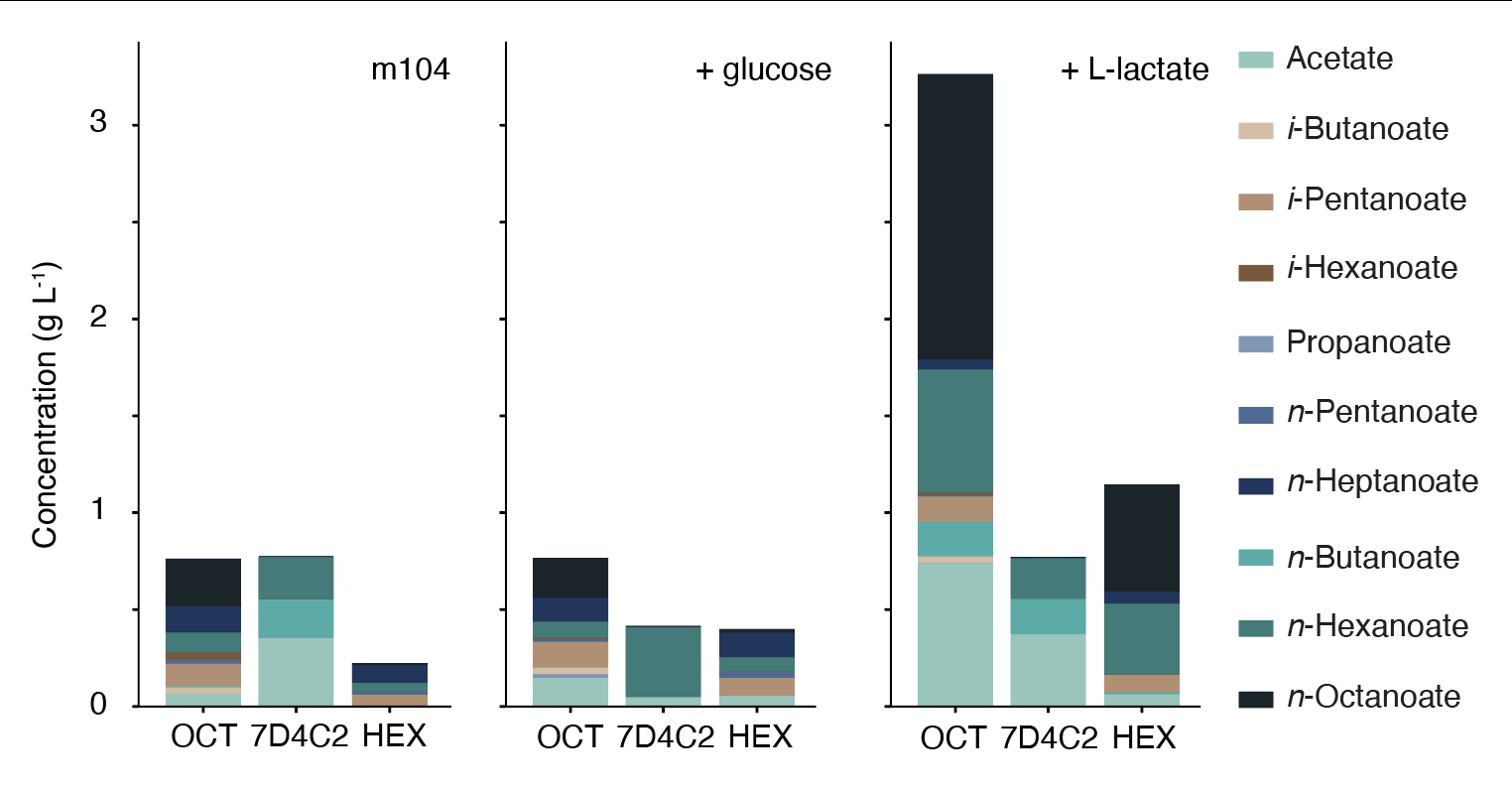

**Figure S7**. Metabolite profiles of *P. octanoica (*OCT), *C. fermentans* (7D4C2), and *M. hexanoica* (MHE) in various m104 media compositions. Added electron donors are labeled in the top corner where m104 represents the basal media (left panel).

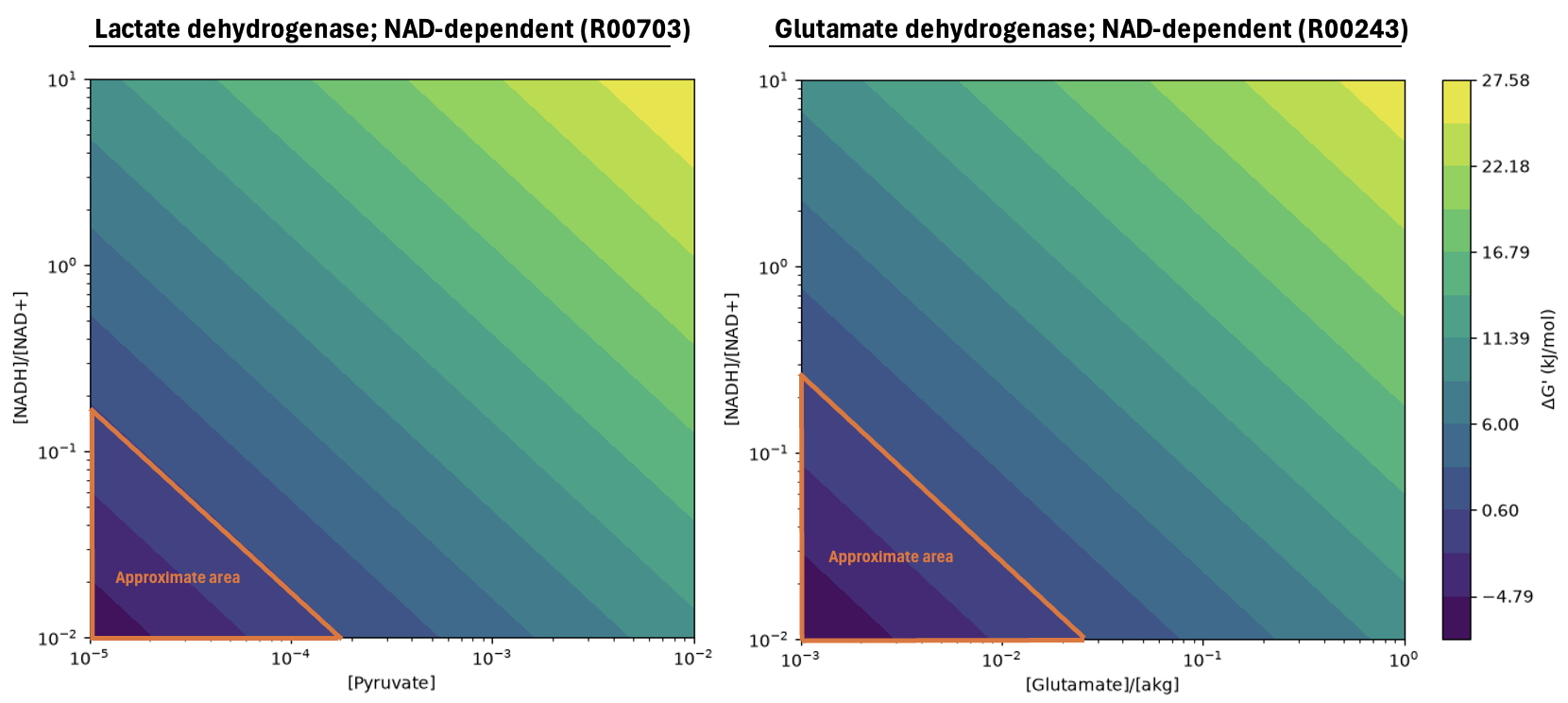

**Figure S8.** Calculated change in Gibb’s free energy for an NAD-dependent lactate dehydrogenase (left) and glutamate dehydrogenase (right) across varying physiological ratios of NADH/NAD^+^, glutamate/α-ketoglutarate, and pyruvate concentrations. The area containing negative values is highlighted. See Table S11 for specific parameters used with eQuilibrator’s API.

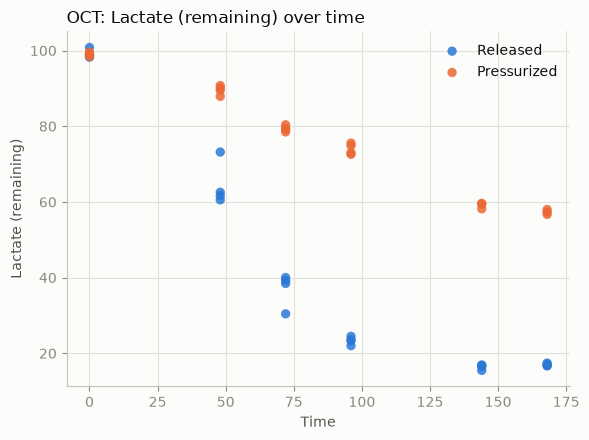

**Figure S9** Lactate utilization by *P. octanoica* cultures grown in 10% m104 with 5 g/L casamino acids under differing partial pressures of H_2_ as simulated by daily release or sparing and re-pressurization with 1 atm of 80:20 H_2_:CO_2_ gas mix.

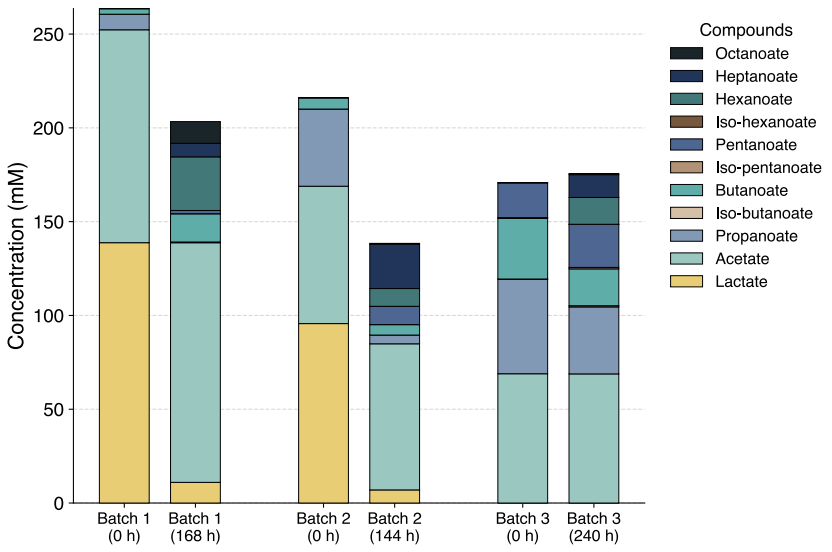

**Figure S10**. Product profile of *P. octanoica* on different batches of SSO (averages of biological replicates shown; n=3, 1, 3, respectively) with varying concentrations of lactate, acetate and propanoate
